# Quantifying interleaflet coupling of phase behavior and observing anti-registered phases in asymmetric lipid bilayers

**DOI:** 10.64898/2026.02.17.706271

**Authors:** Kristen B. Kennison-Cook, Averi M. Cooper, Frederick A. Heberle

**Affiliations:** Department of Chemistry, University of Tennessee, Knoxville, TN 37996, USA; Department of Biochemistry and Biophysics, Stockholm University, Stockholm, Sweden; Science for Life Laboratory, Solna, Sweden

**Keywords:** membrane asymmetry, hemifusion, interleaflet coupling, liquid-liquid phase separation, anti-registered bilayer phases

## Abstract

Model asymmetric lipid bilayers provide a powerful platform for probing how lateral phase behavior in one leaflet is coupled to that of the opposing leaflet. Here, we use calcium-induced hemifusion to generate asymmetric giant unilamellar vesicles (aGUVs) and investigate how lipid composition modulates interleaflet coupling of liquid-liquid phase separation. Symmetric GUVs composed of cholesterol, the high-melting lipid DPPC, and a low-melting phosphatidylcholine (either 14:1-PC or 16:1-PC) were prepared at compositions exhibiting coexisting liquid-ordered (Lo) and liquid-disordered (Ld) phases. Hemifusion with a uniformly mixed supported lipid bilayer composed of the low-melting lipid and cholesterol selectively altered the outer leaflet composition, producing aGUVs with controlled but variable asymmetry. Quantification of outer leaflet exchange using both probe-exit and probe-entry fluorescence measurements revealed substantial vesicle-to-vesicle variability within a given preparation, resulting in overlapping populations of phase-separated and uniformly mixed aGUVs. To account for this variability, we developed a population-based, coupled-distributions framework that enables robust determination of the asymmetric miscibility boundary, defined as the outer leaflet composition at which macroscopic phase separation is suppressed. Independent analyses of probe-exit and probe-entry data yielded consistent boundary locations. Comparing the two lipid systems, we find that aGUVs containing 14:1-PC require significantly greater outer leaflet exchange to abolish phase separation than those containing 16:1-PC. Only in the 14:1-PC system do we observe vesicles exhibiting coexistence of distinct anti-registered phases, a theoretically predicted but rarely observed regime consistent with large hydrophobic mismatch. By expressing both symmetric and asymmetric miscibility boundaries in a common fractional-coordinate framework, we introduce a phenomenological parameter, Δ^∗^, that quantifies the direction and strength of interleaflet coupling of phase behavior. Together, these results demonstrate that modest changes in lipid chain length can markedly alter asymmetric miscibility boundaries and provide a quantitative link between experimental observations, leaflet dominance concepts, and coupled-leaflet theories of membrane organization.

**Statement of Significance:** Membrane asymmetry is a defining feature of eukaryotic cells whose influence on lateral membrane organization remains unclear. Using asymmetric giant vesicles, we find that coexisting liquid-ordered and liquid-disordered domains transition to a uniform appearance as saturated lipid in the outer leaflet is replaced with unsaturated lipid. The extent of exchange required to disrupt phase separation increased with acyl-chain length mismatch, revealing a compositional dependence of interleaflet coupling. In mixtures with greater hydrophobic mismatch, we also observe coexisting anti-registered phases predicted by theory but rarely observed experimentally, providing new constraints for models of coupled-leaflet behavior. By accounting for vesicle-to-vesicle compositional variability, these results provide a framework for measuring asymmetric miscibility boundaries and for connecting asymmetric membrane organization to lipid raft phenomena.

## 1. Introduction

The plasma membrane (PM) separates the cell interior from the outside environment and is centrally involved in many biological processes. In eukaryotic cells, the inner (cytosolic) and outer (exoplasmic) PM leaflets are compositionally asymmetric: the outer leaflet is a mixture of unsaturated phosphatidylcholine (PC) and saturated sphingomyelin (SM) lipids, while the inner leaflet is a mixture of PC, phosphatidylserine, phosphatidylethanolamine, and phosphatidylinositol lipids, nearly all of which are unsaturated (1). The PM of mammalian cells also contains 30-50 mol% cholesterol (2) which, unlike phospholipids, can rapidly transit between leaflets (3-6). While the transbilayer distribution of cholesterol has proven difficult to determine (7), recent measurements indicate that the majority resides in the outer leaflet (8,9).

The striking compositional difference between the inner and outer leaflets raises important questions about lipid miscibility in the PM. Diverse in vitro and in vivo evidence suggests that the outer leaflet, having substantial amounts of both ordered and disordered lipids in addition to cholesterol, may be primed for liquid-liquid phase separation (10). The propensity of outer leaflet lipids to demix dovetails with the membrane raft hypothesis, which proposes a key biological role for dynamic, nanoscale lipid clustering in processes such as signal transduction, membrane trafficking, and viral entry and exit (11,12). In contrast to the outer leaflet, most inner leaflet lipids are unsaturated and fully fluid at physiological temperatures, showing little or no tendency toward phase separation in model membrane studies (13,14). A key question is whether microdomain formation in the PM outer leaflet can induce segregation of inner leaflet lipids and thereby orchestrate the spatial organization of cytosolic proteins that are anchored to the inner leaflet via lipidation (15,16). Equally intriguing is the possibility that the inner leaflet may suppress domain formation in the outer leaflet which, like a loaded spring, would serve as a store of potential energy in the form of unfavorable nearest neighbor interactions. In this case, subtle changes in the composition of either leaflet—for example, induced by the activation of scramblases—could act like a switch that triggers lipid and protein reorganization in both leaflets (17).

Each of the scenarios just described—i.e., dominance of the phase-separated leaflet vs. dominance of the uniform leaflet—has been observed experimentally for specific leaflet compositions in model membranes, as has independent (uncoupled) leaflet phase behavior (18-28). This diverse range of outcomes in biologically relevant lipid mixtures demonstrates a remarkable sensitivity of membrane phase behavior to interleaflet interactions. Building from these observations, theoretical work has identified the surface tension at the midplane interface of a mismatched or “anti-registered” domain (i.e., one characterized by a large difference in order between the two leaflets) as a key parameter that, together with the in-plane line tension, controls the interleaflet coupling of phase behavior and the location of any phase boundaries within the asymmetric composition space (29-33). The magnitude of this surface tension has been estimated from molecular mean-field theory (30,34) and heuristic arguments (35), and measured in experiments (36,37) and simulations (38,39), with the resulting values spanning two orders of magnitude. There remains a need for systematic studies to establish the molecular determinants of interleaflet coupling. Such work relies on the ability to determine the locations of phase boundaries within the asymmetric composition space with relatively high precision, and thus requires robust methods for preparing asymmetric bilayers where the composition of each leaflet can be precisely controlled.

To this end, several techniques for preparing asymmetric vesicles in a range of sizes have been described in the literature (recently reviewed in (40-43)). Methods for preparing asymmetric giant unilamellar vesicles (aGUVs) are particularly valuable for optical microscopy studies of phase behavior and include variations of cyclodextrin-mediated lipid exchange (23,44), water-oil phase transfer (45), and hemifusion (26). The latter of these uses millimolar calcium concentrations to induce hemifusion between symmetric GUVs (sGUVs) and a supported lipid bilayer (SLB), thus facilitating exchange of their outer leaflet lipids (Ca^2+^ is removed by EDTA in the final step). Different fluorescent lipid probes incorporated in the sGUV and SLB are used both to visualize the resulting aGUVs and to quantify the extent of outer leaflet exchange. While the number of studies utilizing calcium-induced hemifusion is still small (26,27,37,46-48), it is notable that all published data show wide variability in lipid asymmetry within a population of aGUVs, often ranging from very little to nearly complete outer leaflet exchange in a single preparation. Although fluorescence-based estimates of exchange for individual vesicles are subject to substantial uncertainty, the observed variability also reflects real vesicle-to-vesicle differences in lipid exchange. As a serendipitous consequence, the degree of asymmetry emerges as a “built-in” variable that can be exploited to gain new insights.

This built-in variability has already proven informative. For example, a recent study from our group investigated the behavior of aGUVs formed by the hemifusion of phase-separated DPPC/DOPC GUVs to an SLB composed of DOPC (37). Sorting the aGUVs in order of increasing outer leaflet exchange revealed a phase boundary at ≈ 60% exchange: beyond this threshold, most aGUVs were visually uniform in fluorescence images, suggesting an abrupt lateral lipid reorganization arising from interactions between the leaflets. The same study estimated a relative uncertainty of ±12-18% in the outer leaflet exchange fraction calculated for individual aGUVs, a surprising result given the considerably smaller uncertainties reported in previous studies. These results highlight both the importance of interleaflet coupling in raft phenomena and the urgent need to understand the sources of error that contribute to compositional uncertainty in aGUVs prepared by hemifusion.

Here, we extend our previous aGUV studies to cholesterol-containing bilayers that more closely mimic the compositional asymmetry of mammalian plasma membranes. We prepared GUVs composed of cholesterol, DPPC, and either 16:1-PC or 14:1-PC as a low-melting lipid—mixtures that exhibit liquid-disordered (Ld) + liquid-ordered (Lo) phase separation in symmetric membranes at 22 °C. Asymmetry was introduced by hemifusion to SLBs composed of the low-melting lipid and cholesterol (80/20 mol%), a composition that is uniformly mixed and enriched in unsaturated lipid relative to the initial symmetric GUV, resulting in aGUV outer leaflets of variable composition. In both systems, we observe populations of aGUVs that are either phase-separated or uniformly mixed, with uniform vesicles exhibiting greater outer leaflet exchange on average, consistent with the presence of a miscibility boundary in asymmetric composition space. To quantitatively determine the location of this boundary, we developed a coupled-distributions framework that accounts for substantial vesicle-to-vesicle variability in outer leaflet composition inferred from fluorescence probe exchange. Rather than treating this variability as experimental noise, the framework explicitly incorporates both genuine heterogeneity in leaflet exchange and compositional uncertainty arising from fluorescence-based measurements, enabling all observed vesicles to contribute information about the underlying exchange distribution and asymmetric miscibility boundary. Using this approach, we find a pronounced shift in the boundary location for aGUVs containing 14:1-PC compared to 16:1-PC, demonstrating that modest changes in acyl chain length—and thus thickness mismatch—strongly influence interleaflet coupling of phase behavior. More broadly, our results highlight that compositional uncertainty is an intrinsic feature of aGUV preparations and must be accounted for when relating asymmetric membrane composition to lateral phase organization.

## 2. Materials and Methods

### 2.1 Materials

Phospholipids 1,2-dipalmitoleoyl-*sn*-glycero-3-phosphocholine (16:1-PC), 1,2-dimyristoleoyl-*sn*-glycero-3-phosphocholine (14:1-PC), and 1,2-dipalmitoyl-*sn*-glycero-3-phosphocholine (DPPC) were purchased from Avanti Polar Lipids (Alabaster, AL) as a dry powder and used as supplied. Cholesterol (Chol) was purchased from Nu-Chek Prep (Elysian, MN) as a dry powder and used as supplied. Phospholipid and cholesterol stock solutions in HPLC-grade chloroform were prepared gravimetrically using an analytical balance with 0.1 mg precision. Fluorescent dyes were purchased from Avanti Polar Lipids as liquid stocks in chloroform and used as supplied: 1-palmitoyl-2-(dipyrrometheneboron difluoride)undecanoyl-*sn*-glycero-3-phosphocholine (TopFluor-PC or TFPC), 1,2-dioleoyl-*sn*-glycero-3-phosphoethanolamine-N-(lissamine rhodamine B sulfonyl) (ammonium salt) (Lissamine Rhodamine-PE or LRPE), and 1,2-dioleoyl-*sn*-glycero-3-phosphoethanolamine-N-(TopFluor AF594) (ammonium salt) (AF594). Dye stock solution concentrations were determined from absorbance measurements: TFPC was measured at 495 nm using an extinction coefficient of 96,900 M^-1^cm^-1^; LRPE was measured at 560 nm with an extinction coefficient of 95,000 M^-1^cm^-1^; and AF594 was measured at 590 nm with an extinction coefficient of 92,000 M^-1^cm^-1^. All stock solutions were stored at −20 °C until use. Sucrose was purchased from Sigma-Aldrich (St. Louis, MO) and dissolved in ultrapure water to make a 100 mM stock solution. Sodium hydroxide was purchased from ThermoFisher Scientific (Waltham, MA) and used to make a 1 M cleaning solution. Sodium chloride was purchased from Sigma-Aldrich. 4-(2-hydroxyethyl)-1-piperazineethanesulfonic acid (HEPES), calcium chloride, and ethylenediaminetetraacetic acid (EDTA) were purchased from ThermoFisher Scientific and used to make various buffer solutions.

### 2.2 Preparation of symmetric GUVs

Chloroform mixtures of lipids were prepared in glass culture tubes using a syringe and repeating dispenser (Hamilton USA, Reno, NV). Samples received a determined volume from chloroform lipid stocks to produce a composition of DPPC/16:1-PC/Chol or DPPC/14:1-PC/Chol = 39/39/22 mol% (≈250 nmol total lipid). Fluorescent probes were added to achieve probe:lipid mol ratios of 1:1000 for TFPC, 1:5000 for LRPE, or 1:2500 for AF594. In probe-exit experiments, TFPC was initially incorporated into the GUV membrane and LRPE served as the red probe in the SLB, whereas in probe-entry experiments, the red probe (LRPE or AF594) was incorporated into the GUV membrane and TFPC was included in the SLB. AF594 was used as a substitute for LRPE in a subset of probe-entry experiments for the DPPC/16:1-PC/Chol system. The concentration of red dyes was kept low to minimize energy transfer from TFPC.

GUVs were prepared by electroformation (49). Briefly, the chloroform solution of lipids and dye was deposited onto the ends of two ITO-covered slides (Delta Technologies, Stillwater, MN). After evaporation of bulk chloroform, the slides were placed in a heated desiccator for 2 h at 55 °C to remove residual solvent, resulting in a dry lipid film coating one end of the slides. A large O-ring was then pressed firmly onto the lipid film on one of the slides to create a chamber, and two small O-rings were placed on the clean end of the slide to act as spacers. After warm sucrose was added to the chamber, the second ITO slide was placed on top, and the assembly was stabilized with a clip. The slide “sandwich” was then placed into a pre-warmed metal block. A positive lead was attached to one slide and a negative lead to the other to create a capacitor. GUVs were produced by applying a 2 Vpp, 10 Hz sine wave for 2 h at 55 °C. The block holding the sandwich was then allowed to cool to 23 °C over 12 h. The GUVs were harvested and subsequently imaged with confocal fluorescence microscopy (CFM) as described below.

### 2.3 Preparation of asymmetric GUVs

#### 2.3.1 SLB formation

SLBs were prepared using previously described protocols with minor modifications (26,50). In brief, chloroform mixtures of lipids were prepared in glass culture tubes using a syringe and repeating dispenser. Samples received a determined volume from chloroform lipid stocks to produce a composition of 16:1-PC or 14:1-PC/Chol = 80/20 mol% (600 nmol total lipid). LRPE or TFPC was added to achieve a probe:lipid mol ratio of 1:10000 or 1:1000, respectively. After adding 600 μL aqueous buffer (1 mM EDTA, 25 mM HEPES, 35 mM NaCl), the tube was mounted on a rapid solvent exchange (RSE) device. RSE exposes the suspension to vacuum while vortexing, thus removing chloroform and producing aqueous vesicles with a broad size distribution and low average lamellarity (51-53). The RSE suspension was then sonicated for 40 min in a cup horn sonicator (QSonica, Newtown, CT) at 50% power to produce small unilamellar vesicles (SUVs) of ≈ 75-100 nm diameter as verified by dynamic light scattering using a Litesizer 100 (Anton Parr, Graz, Austria). After sonication, the SUV solution was further diluted with 2.4 mL buffer to a final concentration of 0.2 mM.

SLBs were prepared in 4-well chambered cover glass (ibidi, Gräfelfing, Germany) that was first cleaned with 1 M NaOH, copiously rinsed with ultrapure water, dried thoroughly at 55 °C under vacuum for 3 min, and dusted with a nitrogen stream. The dish was then plasma cleaned (Harrick Plasma, Ithaca, NY) on the high setting for 2 min 45 s. Immediately after removing the dish from the plasma cleaner, the diluted SUV solution was deposited into the chambered cover glass with ≈ 750 μL pipetted into each well. The SLB dish was left at ambient lab temperature (≈ 22 °C) for 30 min, allowing time for the formation of an SLB as the SUVs adsorbed to the cover glass at the bottom of the chamber. The chamber was then flushed with ultrapure water to remove the remaining vesicles and imaged with CFM to check for quality, ensuring a flat SLB was present with no vesicles stuck to the surface.

#### 2.3.2 Hemifusion

aGUVs were prepared by calcium-induced hemifusion of the GUVs to the SLB, which joins their distal leaflets and allows for lipid exchange via diffusion (26). GUVs were added to a well of the chambered cover glass that contained buffer (35 mM NaCl and 25 mM HEPES, pH 7.4) and the SLB. After allowing ≈ 15 min for the GUVs to settle toward the SLB, a calcium-containing buffer (20 mM CaCl_2_, 25 mM HEPES, 5 mM NaCl, pH 7.4) was added for a final concentration of 5-6 mM CaCl_2_. After 15-30 min, an EDTA-containing buffer (30 mM EDTA, 25 mM HEPES, 5 mM NaCl, pH 7.4) was added to a final concentration of 6-7 mM EDTA in the well to chelate the calcium and terminate hemifusion. We caution that it is often necessary to adjust buffer concentrations to match osmolarities and thus prevent bursting of GUVs. Buffer osmolarities were checked using an OsmoTECH osmometer (Advanced Instruments, Norwood, MA) and adjusted as needed. After osmolarity matching, the EDTA-containing buffer typically had a diluted concentration of ≈ 20-25 mM EDTA. Lastly, the aGUVs were detached from the SLB by gentle pipet mixing with a large-orifice pipette tip.

One goal of our study was to assess the uncertainty in outer leaflet exchange fraction calculated from fluorescence intensity measurements. We performed the hemifusion experiment in two ways with the 16:1-PC mixtures, demonstrated schematically in Fig. 1. In the “probe-exit” experiment, GUVs containing TFPC were hemifused to an SLB to form aGUVs, and the amount of outer leaflet exchange was calculated from the decrease in TFPC fluorescence (Fig. 1a). In the “probe-entry” experiment, GUVs were hemifused to a TFPC-containing SLB and the amount of exchange was calculated from the increase in TFPC fluorescence in the aGUV (Fig. 1b). In both cases, a trace amount of red fluorophore was included in either the SLB (for probe-exit) or GUV (for probe-entry) to aid in visualization. Because we used a very low concentration of the red probe to avoid fluorescence crosstalk, we did not calculate outer leaflet exchange fraction using the red signal. The calculations of exchange fraction using the TFPC signal are described in detail below. In the case of the 14:1-PC mixtures, only the probe-entry experiment was performed.

**Figure 1.**
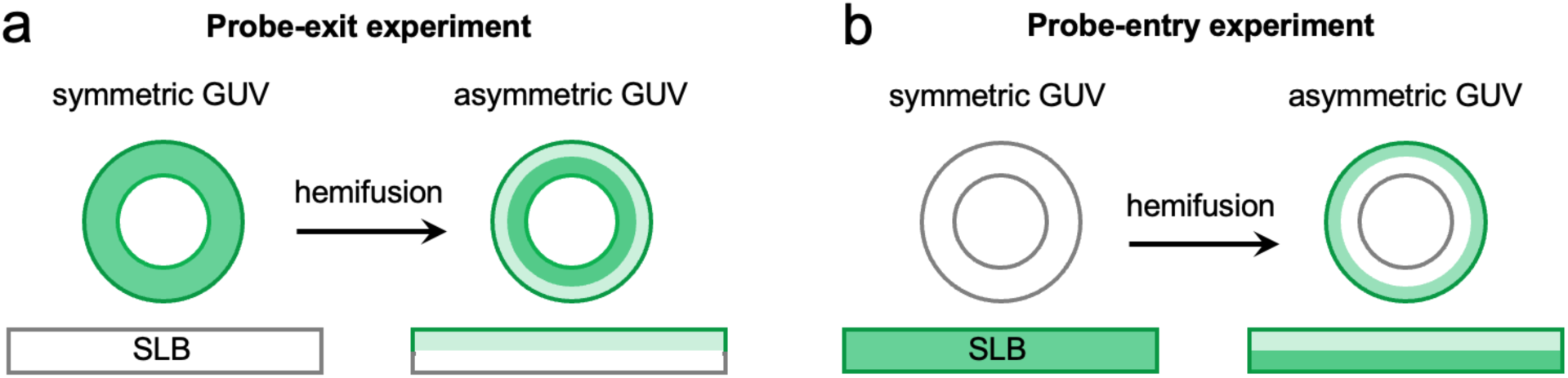
Schematic illustration of hemifusion experiments in this study. (a) In the probe-exit experiment, the fluorescent lipid probe used to quantify exchange is initially present in the GUV. During hemifusion to an SLB that does not contain the probe, the intensity of the GUV decreases. (b) In the probe-entry experiment, the fluorescent lipid is initially present only in the SLB. During hemifusion, intensity thus increases in the asymmetric GUV.

### 2.4 Imaging vesicles

All imaging was performed at 22 °C, maintained with an objective cooling collar (Bioptechs, Butler, PA), using a Nikon C2+ point scanning system attached to a Nikon Eclipse Ti2-E microscope equipped with a Plan Apo Lambda 60×/1.4 NA oil immersion objective (Nikon Instruments, Melville, NY). We first imaged the symmetric GUVs in a well that contained an SLB and ≈ 700 μL of buffer solution (≈ 25 mM HEPES, 35 mM NaCl, pH 7.4) to match the imaging conditions for aGUVs. Approximately 6 μL of the GUV solution was added to the well and allowed to settle for 10 min. We looked for isolated, large, and round unilamellar vesicles to image. Once a suitable GUV was located, we centered it on the screen and set the zoom to 3×. The 488 nm laser was used to image GUVs containing TFPC while the 561 nm laser was used to image GUVs containing LRPE or AF594. For each image, the gain (HV) was set to 90 and the laser power was set to 7. Quarter waveplates (ThorLabs, Newton, NJ) were inserted in the excitation paths to correct for polarization artifacts as described previously (54).

The procedure for imaging asymmetric GUVs was similar to that described above for symmetric GUVs with the following modifications. After the addition of EDTA-containing buffer and gentle pipette mixing to shear aGUVs from the SLB, vesicles were allowed to settle for 10 min. We used a macro to take sequential images in the red (561 nm laser, 546 nm quarter waveplate) and green (488 nm laser, 488 nm quarter waveplate) channels. The observation of fluorescence in both channels allowed us to determine whether a given GUV had exchanged lipid with the SLB, as well as to identify anti-registered phases.

### 2.5 Image analysis to determine the fraction of outer leaflet exchange

All images were saved as .nd2 files and subsequently processed in Fiji (55). To quantify lipid exchange, we analyzed only the green channel of GUVs imaged at the equator. The fluorescence intensity was quantified in the same way for sGUVs prior to hemifusion and aGUVs after hemifusion. First, a circle was drawn on each GUV and the fluorescence intensity profile was extracted radially in 1° angular increments around the equator, yielding 360 values per vesicle. For each angular sector, the maximum pixel intensity was recorded. The equatorial fluorescence, *F*, was then calculated as the average of these 360 peak intensities. We denote this quantity as *F*_*A*_ for asymmetric GUVs and *F*_*S*_ for symmetric GUVs, noting that in all cases *F* represents an averaged vesicle brightness, not an integrated signal. The fraction of exchanged outer leaflet lipids, *ɛ*_*obs*_, for an individual aGUV was then calculated by comparing *F*_*A*_ to the average fluorescence of a population of sGUVs, *F̄*_*S*_. For probe-exit experiments (i.e., where TFPC initially contained in sGUVs diffused into the SLB during hemifusion), *ɛ*_*obs*_ is given by:

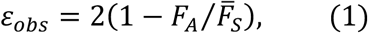

where *F̄*_*S*_ is the average fluorescence intensity of symmetric vesicles measured prior to hemifusion (typically calculated from ∼50 vesicles). For probe-entry experiments (i.e., where TFPC initially contained in the SLB diffused into GUVs during hemifusion), the fraction of outer leaflet exchange is given by:

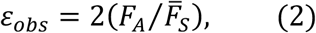

where *F̄*_*S*_ is the average fluorescence intensity of symmetric GUVs (n ≈ 50) containing TFPC at a concentration identical to that of the SLB.

### 2.6 Exchange data analysis

#### 2.6.1 Coupled-distributions model

Exchange data were analyzed with a coupled-distributions model implemented in Mathematica v14.0 (Wolfram Research Inc., Champaign, IL) and described here. Hemifusion produces aGUVs of highly variable composition, ranging from no outer leaflet exchange to nearly complete exchange (26,27,37). Moreover, because hemifusion initiates stochastically in individual vesicles and proceeds over finite time, the experimentally observed ensemble represents a snapshot of a dynamically evolving distribution of exchange fractions. To account for this variability, we use a phenomenological modified exponential distribution that provides a minimal one-parameter description of this skewed ensemble, thus allowing us to quantify the probability of obtaining an aGUV with true exchange fraction *ɛ* ∈ [0,1],

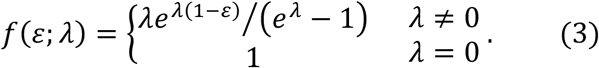

The parameter *λ* ∈ ℝ skews the distribution toward smaller (*λ* > 0) or larger (*λ* < 0) values of *ɛ*, with the special case *λ* = 0 corresponding to a uniform distribution in which all values of *ɛ* are equally likely.

The model must also account for the uncertainty in estimating *ɛ* from fluorescence intensity measurements that are subject to error. We introduce the notation *ɛ*_*obs*_ to distinguish the observed exchange fraction of an aGUV (i.e., calculated from fluorescence measurements) from its true exchange fraction *ɛ*. Error propagation analysis of Eqs. 1 and 2 reveals that uncertainty in *ɛ*_*obs*_ has a functional dependence on *ɛ* that is different for the probe-exit and probe-entry experiments (Supporting Information S5). Accordingly, we use normal distributions *g*_*exit*_ and *g*_*entry*_ to model the probability that an aGUV with true exchange value *ɛ* will produce a calculated exchange value *ɛ*_*obs*_, where the width of the distribution depends on *ɛ* according to Eq. S8a (for probe-exit) or Eq. S8b (for probe-entry):

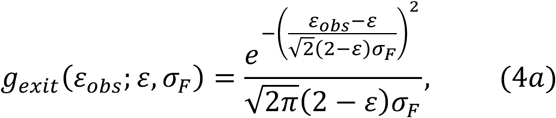

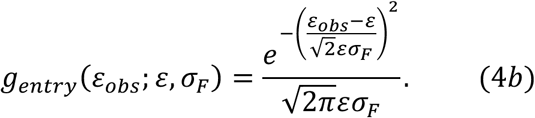

In Eqs. 4, *σ*_*F*_ is a composite uncertainty that accounts for all sources of error inherent to measuring the average fluorescence intensity of a GUV from an equatorial confocal image.

The final addition to the model is the inclusion of a phase boundary in asymmetric composition space, as predicted by coupled-leaflet theories (29,32). We denote the location of this boundary along the exchange coordinate by *ɛ*^∗^, corresponding to the value of the outer-leaflet exchange fraction at which macroscopic phase separation becomes thermodynamically unfavorable. In this framework, asymmetric vesicles with *ɛ* < *ɛ*^∗^ are phase-separated, whereas asymmetric vesicles with *ɛ* > *ɛ*^∗^ are laterally uniform within each leaflet. When hemifusion generates a distribution of *ɛ* values that straddles *ɛ*^∗^, both phenotypes will naturally be present in the population. Combining Eqs. 3-4, the probability that a phase-separated aGUV (i.e., for which 0 < *ɛ* < *ɛ*^∗^) will result in a measured exchange fraction 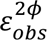 is

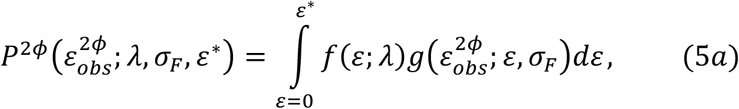

while the probability that a uniform aGUV (i.e., for which *ɛ*^∗^ < *ɛ* < 1) will result in a measured exchange fraction 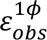 is

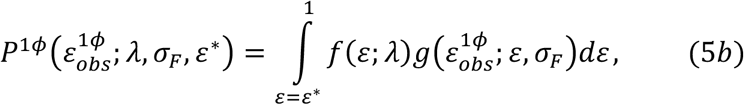

with *g* replaced by *g*_*exit*_(Eq. 4a) or *g*_*entry*_(Eq. 4b) for a probe-exit or probe-entry experiment, respectively. The probability distributions in Eqs. 5 are coupled by the shared parameters *ɛ*^∗^, *λ*, and *σ*_*F*_, and are thus refined jointly in a global fit. The integrals do not have closed form solutions but can be numerically evaluated for data fitting.

#### 2.6.2 Data fitting

We denote the set of experimentally observed exchange fractions 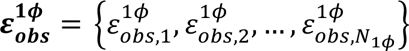 and 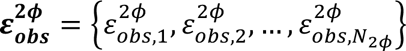 for uniform and phase-separated aGUVs, respectively. The exchange datasets were binned and normalized to produce probability density histograms in which the total probability of all observations (i.e., uniform and phase-separated) was unity. The normalized uniform and phase-separated histograms were then jointly analyzed by fitting to Eqs. 5 using Mathematica’s NonlinearLeastSquares function with default settings. The adjustable parameters were *ɛ*^∗^, *λ*, and *σ*_*F*_.

## 3. Results

### 3.1 Generating asymmetric GUVs by hemifusion produces a spectrum of phase phenotypes

Asymmetric giant unilamellar vesicles (aGUVs) were prepared by calcium-induced hemifusion between symmetric GUVs and supported lipid bilayers (SLBs), a process that joins the outer leaflet of the GUV to the distal leaflet of the SLB and enables lipid exchange by lateral diffusion (Fig. 1). Removal of calcium terminates hemifusion, yielding aGUVs whose inner and outer leaflet compositions generally differ from one another. Because lipid exchange during hemifusion is diffusion-limited and vesicle-specific, a single preparation produces a heterogeneous population of aGUVs that span a wide range of leaflet asymmetries (26,27,37,47,48).

A direct consequence of this heterogeneity is that aGUVs within the same preparation can exhibit markedly different phase behavior. Some vesicles retain coexisting liquid-ordered (Lo) and liquid-disordered (Ld) domains similar to their symmetric precursors, whereas others appear uniformly mixed at the same temperature. In addition to these cases, we occasionally observe vesicles with more complex fluorescence patterns such as spatially modulated or anti-registered domains. Thus, hemifusion generates a spectrum of membrane phase phenotypes rather than a single, well-defined asymmetric composition.

In this study, we examined aGUVs prepared from two ternary lipid mixtures that differ in the chain length of the low-melting phospholipid. In each of these mixtures, several hundred hemifused aGUVs were analyzed, revealing a rich variety of phase phenotypes reproducibly across independent preparations. While the quantitative consequences of this chemical difference are analyzed below, the qualitative features described here—heterogeneous leaflet exchange and the coexistence of phase-separated and uniform vesicles—are common to both systems. These observations motivate the need for a quantitative measure of leaflet exchange in individual aGUVs and for a robust framework to relate that exchange to membrane phase behavior, as described in the following sections.

### 3.2 Quantifying outer-leaflet exchange in individual aGUVs

To relate the diverse phenotypes described above to leaflet composition, we quantified the extent of outer-leaflet lipid exchange for individual aGUVs using fluorescence intensity measurements as previously described (26). In hemifusion experiments, exchange was inferred either from the loss of a fluorescent probe initially present in the GUV (“probe-exit” experiments) or from the gain of probe initially present in the SLB (“probe-entry” experiments) (Fig. 1). In both cases, the measured fluorescence intensity in a central confocal slice was used to calculate an apparent exchange fraction, denoted *ɛ*_*obs*_, which represents the fraction of the outer GUV leaflet replaced by SLB lipids. The use of such equatorial fluorescence images to characterize vesicle phase behavior and quantify exchange is supported by confocal *z*-stack measurements showing that Ld phase fractions measured at the equator accurately represent overall vesicle phase fractions (Fig. S10).

Quantification of *ɛ*_*obs*_requires reference measurements on symmetric GUVs prepared under identical conditions, which serve both to establish baseline fluorescence intensities and to characterize vesicle-to-vesicle variability in probe signal. For each lipid mixture and probe configuration, symmetric control GUVs were imaged in parallel with hemifused samples. These controls define the normalization used to calculate *ɛ*_*obs*_in both probe-exit and probe-entry experiments and provide lower-bound estimates of fluorescence variability relevant to exchange measurements. Data from these symmetric control experiments are summarized in Supporting Information Section S2 (Figs. S1-S4 and Tables S1-S3).

In total, *ɛ*_*obs*_was determined for 658 aGUVs across both lipid mixtures using a combination of probe-exit and probe-entry measurements (Tables S4-S6). Representative aGUVs from both lipid mixtures, drawn from the full datasets and arranged in order of increasing *ɛ*_*obs*_, are shown in Fig. 2. Vesicles with small values of *ɛ*_*obs*_typically exhibit coexisting Lo and Ld domains similar to those observed in symmetric GUVs, whereas vesicles with large *ɛ*_*obs*_are predominantly uniform in fluorescence intensity. Intermediate values of *ɛ*_*obs*_are associated with a range of behaviors, including vesicles that remain phase-separated as well as vesicles that appear uniformly mixed. This overlap reflects two distinct effects: first, hemifusion produces a genuine vesicle-to-vesicle variation in the true extent of outer-leaflet exchange; and second, the value of *ɛ*_*obs*_ inferred for any individual aGUV is subject to substantial uncertainty due to the limitations of fluorescence-based measurements (the sources and magnitude of uncertainty associated with *ɛ*_*obs*_are analyzed in detail in the Supporting Information, Section S5). As a result, *ɛ*_*obs*_is not, by itself, a reliable predictor of phase behavior for any single aGUV, as discussed below.

**Figure 2.**
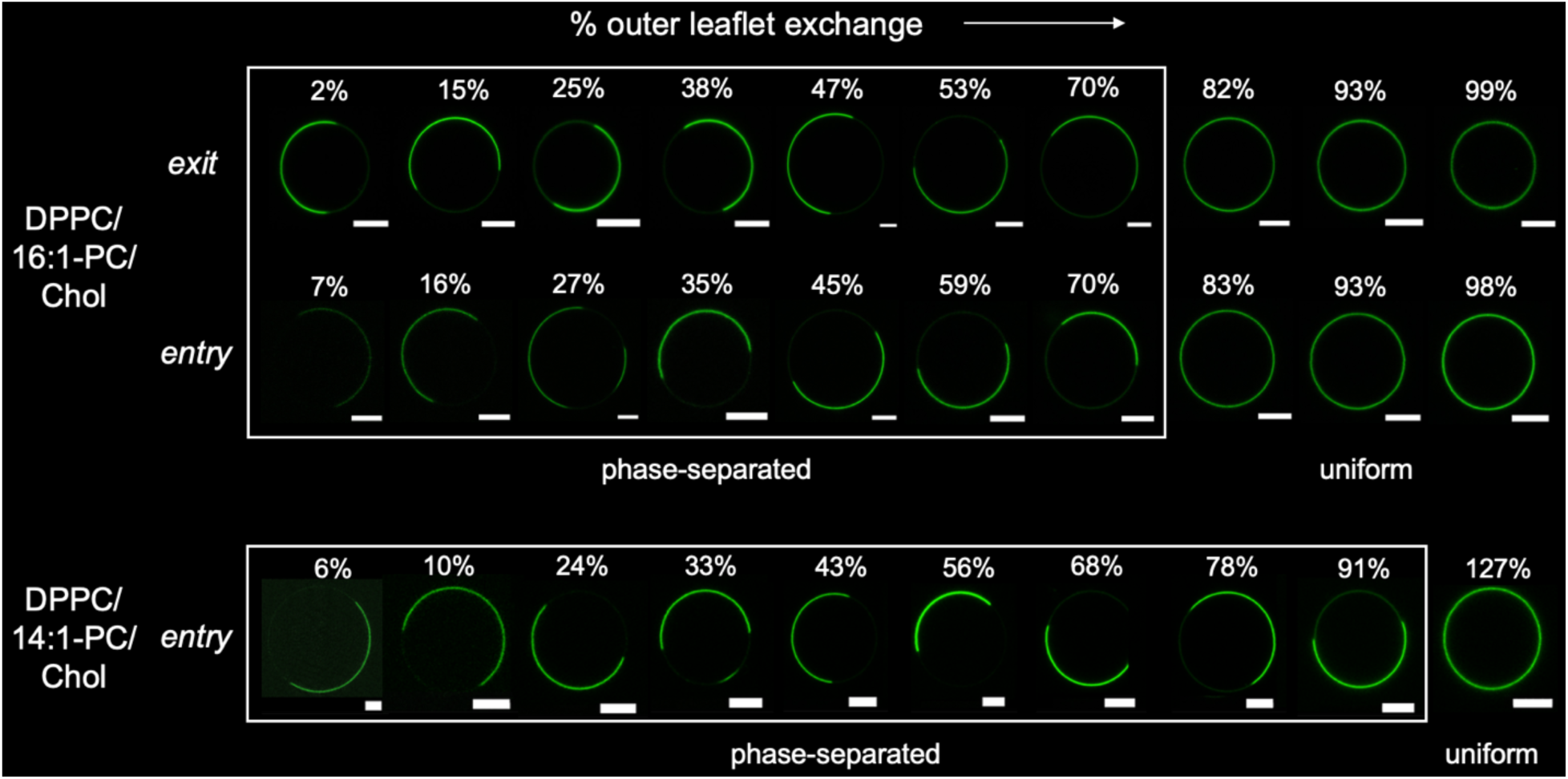
Asymmetric GUVs arranged by apparent outer-leaflet exchange fraction, *ɛ*_*obs*_. Confocal fluorescence images of representative asymmetric GUVs prepared by hemifusion for the DPPC/16:1-PC/Chol and DPPC/14:1-PC/Chol systems, drawn from the full datasets and arranged in order of increasing apparent exchange fraction *ɛ*_*obs*_ as calculated from fluorescence intensity measurements. For small *ɛ*_*obs*_, vesicles typically exhibit coexisting liquid-ordered (Lo) and liquid-disordered (Ld) domains, whereas vesicles with large *ɛ*_*obs*_ are predominantly uniform in fluorescence intensity. Scale bars are 5 µm.

To summarize, the broad distribution of *ɛ*_*obs*_ values observed within a given preparation encodes both real physical heterogeneity in leaflet exchange across the aGUV population and measurement uncertainty associated with quantifying *ɛ*_*obs*_for individual vesicles. While the latter limits the interpretability of *ɛ*_*obs*_at the single-vesicle level, the former provides meaningful information at the population level, thus motivating a statistical treatment of *ɛ*_*obs*_distributions to determine asymmetric miscibility boundaries, as described in the following section.

### 3.3 Inferring asymmetric miscibility boundaries from heterogeneous aGUV populations

Figure 3a shows the full distributions of *ɛ*_*obs*_for all analyzed aGUVs from both lipid mixtures, broken down by lipid system, mode of exchange measurement, and observed phase phenotype (i.e., uniform, two-phase, or three-phase; see Tables S4-S6 for vesicle counts by phenotype and condition). Darker shades within the two-phase distributions indicate vesicles exhibiting modulated phase patterns, examples of which are shown in Fig. S6. It is clear from Fig. 3a that aGUVs prepared under nominally identical conditions exhibit both phase-separated and uniform phenotypes over overlapping ranges of *ɛ*_*obs*_. This overlap precludes assigning a miscibility boundary based on a single threshold value of *ɛ*_*obs*_ for individual vesicles. Instead, determination of a phase boundary requires analysis of the full population of aGUVs while explicitly accounting for vesicle-to-vesicle variability in leaflet exchange and measurement uncertainty.

**Figure 3.**
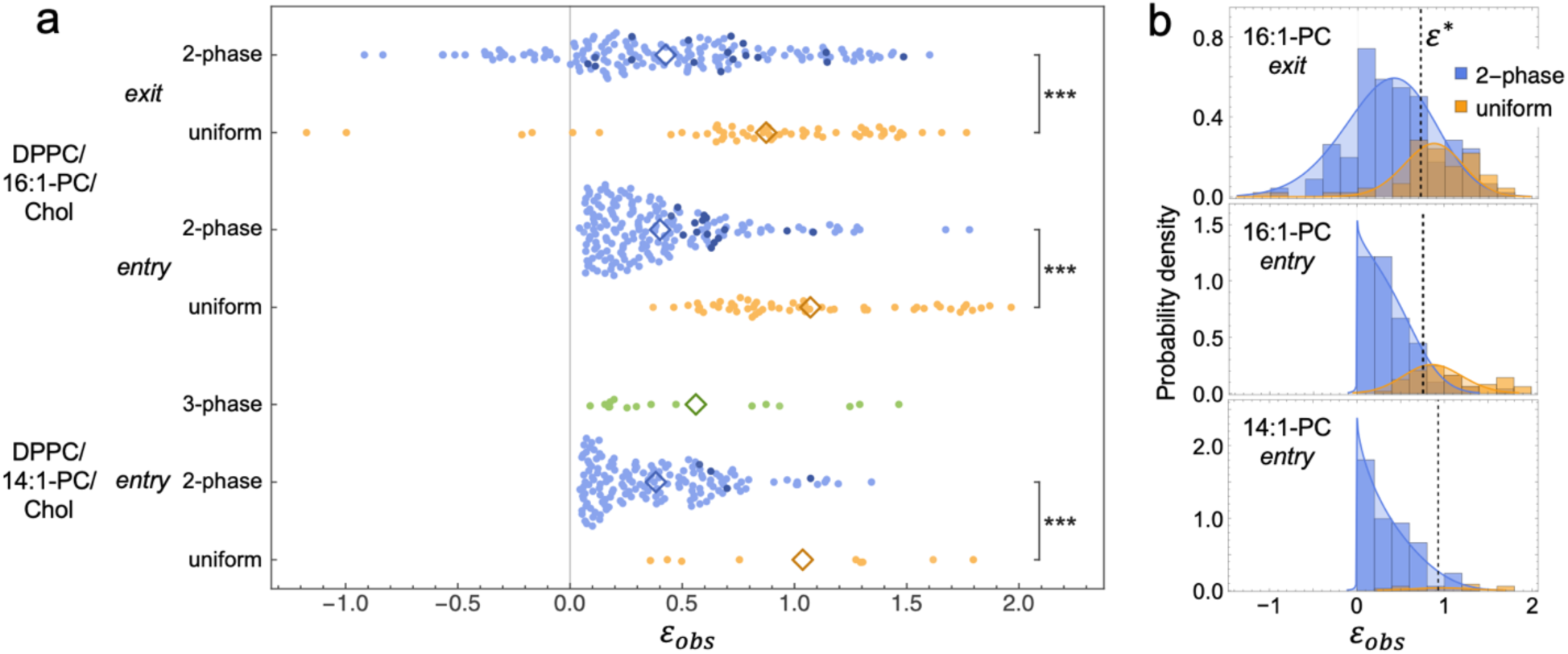
Population-based determination of the asymmetric miscibility boundary. (a) Distributions of the apparent outer-leaflet exchange fraction, *ɛ*_*obs*_, for aGUVs obtained from DPPC/16:1-PC/Chol (probe-exit and probe-entry modes) and DPPC/14:1-PC/Chol (probe-entry mode). The different distributions correspond to distinct phase phenotypes: uniform (orange), 2-phase (blue), and 3-phase (green). Within the 2-phase distributions, the darker shade of blue indicates aGUVs exhibiting modulated phase patterns. Open diamonds denote the mean of each distribution. The full range of obtained *ɛ*_*obs*_ is shown, including values outside the physical range (i.e., *ɛ*_*obs*_< 0 or > 1), which arise primarily from fluorescence measurement uncertainty. (b) Histograms of uniform and 2-phase distributions from panel (a) overlaid with the best-fit curves obtained from fitting to the coupled-distributions model. Best-fit parameters are given in Table 1.

**Table 1.** Summary of the coupled-distributions model applied to DPPC/16:1-PC/Chol and DPPC/14:1-PC/Chol aGUVs.

| Parameter | DPPC/16:1-PC/Chol |  |  | DPPC/14:1-PC/Chol |
| --- | --- | --- | --- | --- |
|  | *Probe-exit* | *Probe-entry* | *Global fit* | *Probe-entry* |
| $N_{2\phi}$ | 172 | 193 | | 157 |
| $N_{1\phi}$ | 57 | 54 | | 9 |
| $\varepsilon^*$ | $0.72 \pm 0.05$ | $0.76 \pm 0.03$ | $0.75 \pm 0.02$ | $0.93 \pm 0.04$ |
| $\sigma_F$ | $0.29 \pm 0.02$ | $0.38 \pm 0.06$ | $0.31 \pm 0.02$ | $0.38 \pm 0.11$ |
| $\lambda$ | $0.5 \pm 0.5$ | $0.3 \pm 0.3$ | $0.4 \pm 0.3$ | $1.3 \pm 0.4$ |

To this end, we jointly analyzed the *ɛ*_*obs*_ data using a coupled-distributions framework. The results are shown in Fig. 3b, where the *ɛ*_*obs*_ distributions corresponding to uniform and two-phase vesicles from Fig. 3a are replotted as histograms. By fitting the model simultaneously to the two histograms—whose shapes are intrinsically linked through a shared underlying exchange distribution and a common phase boundary location—we determined the asymmetric miscibility boundary, *ɛ*^∗^, defined as the value of the true outer-leaflet exchange fraction corresponding to the transition between phase-separated and uniform regimes. The model explicitly accounts for uncertainty in the fluorescence-based determination of *ɛ*_*obs*_ and allows data from all vesicles— including those with unphysical values of *ɛ*_*obs*_—to contribute to the analysis without post hoc exclusion. Best-fit model curves are shown in Fig. 3a as solid lines, and the resulting values of *ɛ*^∗^for each dataset are indicated by vertical dashed lines. Obtained parameters and associated uncertainties (determined by Monte Carlo resampling of vesicle counts within the coupled-distributions framework) are summarized in Table 1. As discussed below, comparison of *ɛ*^∗^ values between the two lipid mixtures reveals a pronounced dependence of the asymmetric miscibility boundary on lipid chain length.

### 3.4 Asymmetric miscibility boundaries differ between DPPC/16:1-PC/Chol and DPPC/14:1-PC/Chol

To compare asymmetric phase behavior across lipid mixtures, we next examine the values of the asymmetric miscibility boundary, *ɛ*^∗^, obtained from the population-level analysis described above. For the DPPC/16:1-PC/Chol system, independent fits to probe-exit and probe-entry datasets yield statistically consistent values of *ɛ*^∗^, with best-fit values of *ɛ*^∗^= 0.72 ± 0.05 and *ɛ*^∗^ = 0.76 ± 0.03, respectively, indicating that the inferred boundary location is robust to the direction of probe exchange. From a global fit in which *ɛ*^∗^ was constrained to be the same across probe-entry and probe-exit datasets, we obtain *ɛ*^∗^= 0.75 ± 0.02 for DPPC/16:1-PC/Chol. In contrast, analysis of the DPPC/14:1-PC/Chol system—measured in probe-entry mode—yields a significantly larger value of *ɛ*^∗^ = 0.93 ± 0.04, demonstrating that phase separation persists to higher levels of outer-leaflet exchange in this mixture (Table 1). Taken together, these results establish a clear shift in the location of the asymmetric miscibility boundary between the two lipid mixtures.

### 3.5 Context from symmetric bilayers: phase-boundary location along the aGUV exchange trajectory

To interpret how leaflet asymmetry alters phase behavior in aGUVs, it is essential to establish the phase-diagram context of the corresponding fully symmetric ternary mixtures. We therefore determined where macroscopic Ld+Lo phase separation is lost in symmetric bilayers along the specific composition trajectory relevant to our aGUV experiments, i.e., the path that approximates the compositional evolution experienced by the *outer leaflet* of an aGUV during lipid exchange. Figure 4 summarizes measurements performed along this exchange-relevant trajectory, which connects the initial symmetric GUV composition (39/39/22 mol%, *s* = 0) to the SLB composition (low-*T*_*M*_-PC/Chol = 80/20 mol%, *s* = 1) as shown in Fig. 4a. Here, *s* denotes the fractional distance along the exchange-relevant composition trajectory and therefore provides a direct symmetric reference for interpreting aGUV phase behavior.

**Figure 4.**
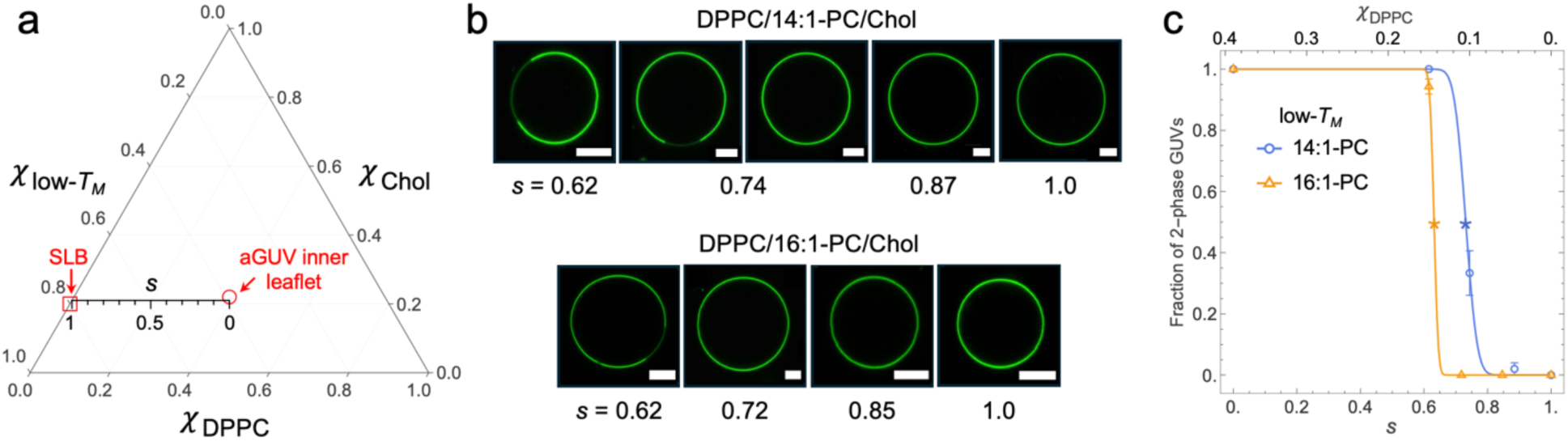
Symmetric phase boundary location along the exchange-relevant composition trajectory. (a) Ternary plot for DPPC/low-*T_M_*-PC/Chol showing the exchange-relevant trajectory used in the hemifusion experiments. The trajectory connects the initial symmetric GUV composition (39/39/22 mol%, red circle, *s* = 0) to the SLB composition (low-*T*_*M*_-PC/Chol = 80/20 mol%, red square, *s* = 1). (b) Representative confocal equatorial images of *symmetric* GUVs prepared along the *s* trajectory for mixtures containing 14:1-PC (top row) or 16:1-PC (bottom row) as the low-*T_M_* lipid. Images illustrate the loss of macroscopic Lo/Ld phase separation as *s* increases. Scale bars, 5 µm. (c) Fraction of symmetric GUVs exhibiting macroscopic phase coexistence as a function of *s* for DPPC/14:1-PC/Chol (blue circles) and DPPC/16:1-PC/Chol (orange triangles). Solid lines show sigmoidal fits used to determine the symmetric miscibility boundary *s*^∗^, defined as the sigmoidal inflection point and indicated by asterisks in panel (c). Counts of uniform and phase-separated symmetric GUVs for each of the examined compositions are given in Table S7 (DPPC/14:1-PC/Chol) and Table S8 (DPPC/16:1-PC/Chol); error bars reflect binomial counting uncertainty. The upper horizontal axis shows the corresponding DPPC mole fraction, *χ*_FGGH_, along the same trajectory.

Symmetric GUVs prepared at the initial composition (*s* = 0) exhibit robust macroscopic Lo+Ld coexistence: large, stable domains are present in every vesicle, with Ld area fractions narrowly distributed around 0.5 (Tables S1-S3). This indicates that the starting composition lies well within the two-phase region for both lipid mixtures and is not close to a miscibility boundary or critical point. As the composition is shifted along the trajectory toward the SLB endpoint, symmetric GUVs eventually lose macroscopic phase separation, revealing a clear transition from vesicles dominated by two-phase coexistence to predominantly uniform vesicles (Fig. 4b). Summaries of these surveys are provided in Tables S7 and S8.

To quantify the location of the symmetric miscibility boundary, we analyzed the GUV data by fitting the fraction of phase-separated vesicles as a function of *s* to a sigmoidal function (Fig. 4c). From these fits, we define *s*^∗^(the inflection point of the fitted curve) as the fractional distance along the trajectory at which symmetric GUVs transition from predominantly phase-separated to predominantly uniform. For the DPPC/16:1-PC/Chol system, this analysis yields *s*^∗^ = 0.63 ± 0.01, while for DPPC/14:1-PC/Chol we obtain *s*^∗^ = 0.73 ± 0.03. These values provide a quantitative measure of the phase boundary location in fully symmetric GUVs, expressed in the same fractional-coordinate framework used to describe lipid exchange in aGUVs. As such, *s*^∗^ can be directly compared to the phase boundary location *ɛ*^∗^determined from asymmetric GUV experiments to define a phenomenological measure of interleaflet coupling, as discussed below.

### 3.6 A phenomenological measure of interleaflet coupling

The asymmetric and symmetric miscibility boundaries determined above can be combined to define a compact phenomenological measure of interleaflet coupling. Specifically, we define

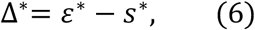

which quantifies the shift of the asymmetric miscibility boundary relative to the corresponding symmetric boundary along the same exchange-relevant composition trajectory.

For both lipid mixtures examined here, Δ^∗^ is positive, indicating that phase separation in asymmetric bilayers persists to larger values of outer-leaflet exchange than would be expected based on symmetric bilayer behavior alone. For DPPC/16:1-PC/Chol, we find *s*^∗^= 0.63 ± 0.01 and *ɛ*^∗^= 0.75 ± 0.02, yielding Δ^∗^ = +0.12 ± 0.02. We find a substantially larger shift for DPPC/14:1-PC/Chol: the corresponding values are *s*^∗^ = 0.73 ± 0.03 and *ɛ*^∗^ = 0.93 ± 0.04, giving Δ^∗^ ≈ +0.20 ± 0.05. These values provide a quantitative basis for comparing the strength of interleaflet coupling between the two systems.

### 3.7 Observation of anti-registered phases in DPPC/14:1-PC/Chol aGUVs

A total of 658 hemifused aGUVs were analyzed across both lipid mixtures examined in this study (DPPC/14:1-PC/Chol and DPPC/16:1-PC/Chol). The overwhelming majority of these vesicles exhibited fluorescence phenotypes broadly consistent with predictions of coupled-leaflet theories, as inferred from equatorial intensity patterns of TFPC and LRPE under the assumption that both probes preferentially partition into the Ld phase. In particular, phase-separated vesicles displayed fluorescence patterns expected for either (i) coexisting ordered and disordered phases with fully registered leaflets, or (ii) coexistence of a fully registered disordered phase with an anti-registered phase consisting of a disordered outer leaflet and ordered inner leaflet. As noted by Williamson and Olmsted, these two cases are difficult to distinguish experimentally because both lie on positively sloped tielines and thus produce similar apparent colocalization of outer- and inner-leaflet probes (56).

Remarkably, 16 vesicles—all from the DPPC/14:1-PC/Chol system—showed clear signatures of anti-registered (AR) phases, revealing a rich phenomenology of asymmetry generation during hemifusion. To describe these cases, we adopt a notation for the physical state of each leaflet: registered phases are denoted R_O_ or R_D_ (indicating ordered or disordered states in both leaflets), whereas anti-registered phases are denoted AR_OD_ or AR_DO_, where the subscript lists, in order, the states of the outer and inner leaflets (i.e., AR_OD_ indicates an ordered outer leaflet and disordered inner leaflet, while AR_DO_ indicates the reverse).

The 16 vesicles with AR phases could be grouped into four categories, representative examples of which are shown in Fig. 5:

1. Two vesicles showed R_O_+R_D_+AR_DO_, which is one of the expected quasi-equilibrium states for an aGUV formed from an initially phase-separated (R_O_+R_D_) sGUV after increasing the fraction of unsaturated lipid in the outer leaflet (Fig. 5a,e). This outcome is fully consistent with theoretical leaflet-leaflet phase diagrams, given the direction of the hemifusion-induced compositional trajectory through asymmetric composition space (e.g., (29,32)).
2. Nine vesicles exhibited coexistence of AR_DO_+AR_OD_+R_O_ (i.e., both AR phases and the registered ordered phase), with R_O_ always occupying a smaller area fraction than in the initially symmetric GUVs (Fig. 5b,f). This outcome can be interpreted as a metastable three-phase coexistence predicted by Williamson and Olmsted (32,56,57).
3. Two aGUVs contained AR_DO_+AR_OD_ with at most a trace amount of R_O_ (Fig. 5c,g).
4. Three vesicles appeared to contain R_O_+R_D_+AR_OD_, a pattern that would be expected if hemifusion resulted in a net gain in saturated lipids in the outer leaflet at the expense of unsaturated lipids (Fig. 5d,h). This is opposite to the exchange occurring in our experiments, in which unsaturated lipids are supplied to the outer leaflet from the SLB, making these cases surprising and suggestive of additional mechanistic complexity.

**Figure 5.**
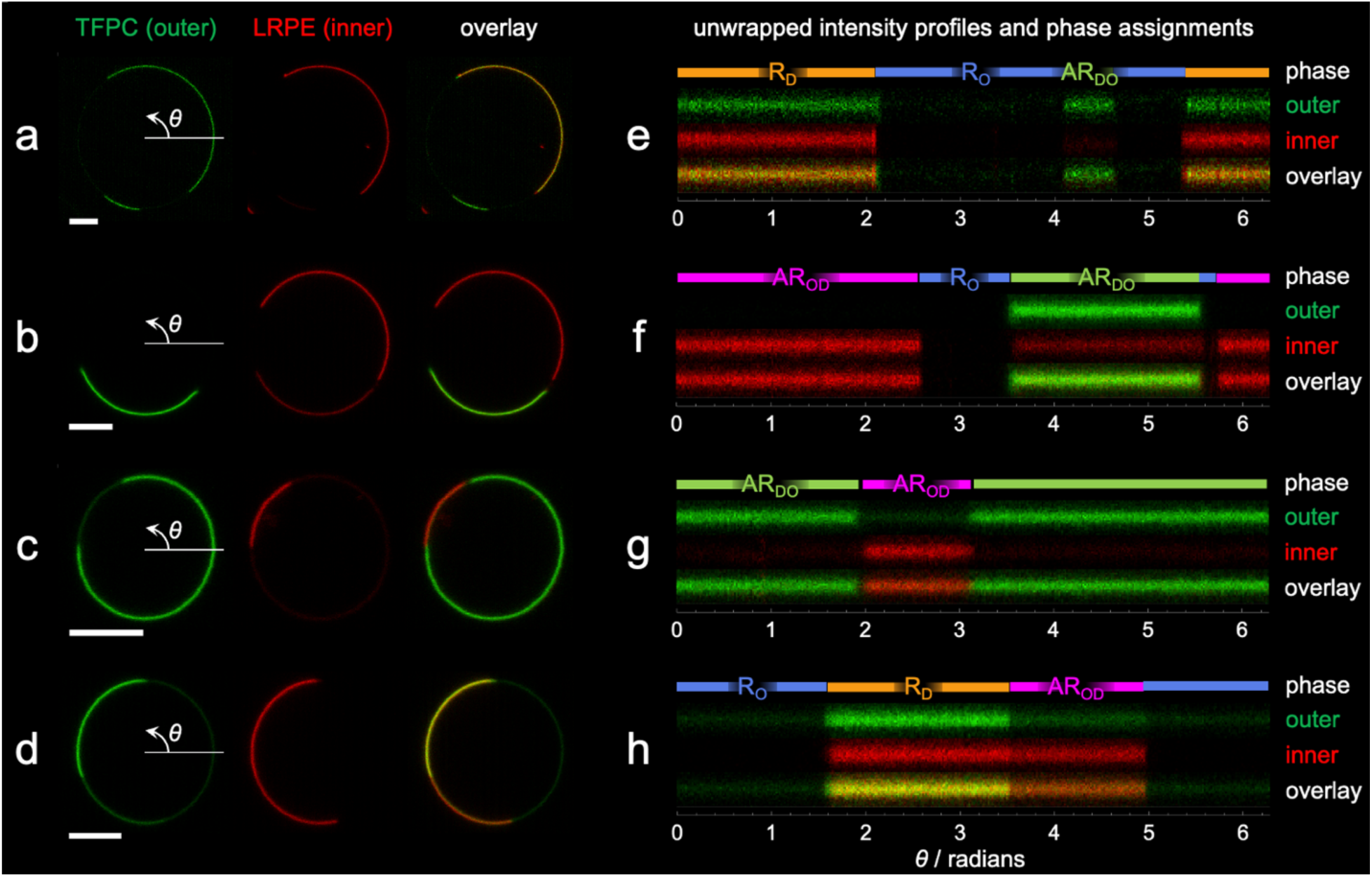
Representative anti-registered (AR) phase phenotypes in DPPC/14:1-PC/Chol asymmetric GUVs. (a–d) Equatorial confocal slices of four representative aGUVs exhibiting AR phase behavior. For each vesicle, fluorescence from the outer leaflet probe TFPC (green), the inner leaflet probe LRPE (red), and their overlay are shown. Arrows indicate the direction of increasing azimuthal angle *θ* used for intensity unwrapping. (e–h) Corresponding unwrapped fluorescence intensity profiles as a function of *θ*, with phase assignments based on the relative intensities of the two probes. Colored bars above each profile indicate registered disordered (R_D_), registered ordered (R_O_), and anti-registered (AR_DO_, AR_OD_) regions. Panels (a,e), (b,f), (c,g), and (d,h) correspond to the same vesicle. Scale bars are 5 µm.

AR phases were observed in the 14:1-PC system but not in the 16:1-PC system, consistent with theoretical predictions that AR phases are favored by greater hydrophobic mismatch between ordered and disordered phases (32,56,57). In addition, the observation of coexisting AR phases (categories 2 and 3 above) is consistent with the presence of multiple AR states predicted in coupled-leaflet phase diagrams. Finally, we note that similar phenotypes were reported in ref. (46), in which some hemifusion-generated aGUVs displayed AR domains coexisting with each other and with registered phases, although the significance of these observations was not discussed.

## 4. Discussion

### 4.1 Quantifying interleaflet coupling through shifts in phase boundaries

Our experimental systems consisted of the initially phase-separated compositions DPPC/16:1-PC/Chol or DPPC/14:1-PC/Chol at 39/39/22 mol % and 22 °C. As established in Section 3.5, these mixtures lie well within the Ld+Lo coexistence region under symmetric conditions. After exchange of outer leaflet lipids by calcium-induced hemifusion, a substantial fraction of the resulting aGUVs in both mixtures exhibited uniform fluorescence, indicating loss of macroscopic phase separation, while others retained clear Lo+Ld coexistence (Fig. 2). For both systems, the average exchanged lipid fraction of uniform aGUVs was significantly larger than that of phase-separated aGUVs (Fig. 3a), consistent with the presence of a miscibility boundary in asymmetric composition space. In both systems, Lo domains in the inner leaflet were progressively destabilized as high-melting DPPC in the outer leaflet was replaced by low-melting lipid (either 16:1-PC or 14:1-PC).

To interpret these observations, it is useful to compare phase boundary locations in symmetric and asymmetric bilayers along the same exchange-relevant composition trajectory. We quantified the location of the miscibility boundary in fully symmetric GUVs by defining *s*^∗^ as the fractional distance along this trajectory at which vesicles transition from predominantly phase-separated to predominantly uniform (Fig. 4a). Independently, we defined *ɛ*^∗^ from the aGUV experiments as the outer leaflet composition at which macroscopic phase separation is lost in asymmetric vesicles (Fig. 3b). While these quantities are defined in different experimental contexts, they are expressed in the same fractional-coordinate framework and can therefore be directly compared, revealing a systematic offset between symmetric and asymmetric phase boundary locations. For DPPC/16:1-PC/Chol, this offset corresponds to Δ^∗^ = +0.12 ± 0.02, whereas for DPPC/14:1-PC/Chol we obtain a substantially larger shift of Δ^∗^ = +0.20 ± 0.05. Thus, in both systems, macroscopic phase separation persists in aGUVs at outer leaflet compositions that would be uniformly mixed in a fully symmetric bilayer, with this effect being roughly twice as strong in the 14:1-PC system as in the 16:1-PC system.

We interpret Δ^∗^as a phenomenological measure of net interleaflet coupling along the exchange-relevant trajectory. In the context of our hemifusion experiments, a positive value of Δ^∗^ indicates that coupling to the inner leaflet stabilizes demixing in the outer leaflet beyond the point at which it would otherwise be lost under symmetric conditions. In this sense, Δ^∗^ quantifies the extent to which one leaflet biases the miscibility of the other, without requiring identification of a specific microscopic origin for that bias. While coupled-leaflet theories attribute such shifts in phase boundary location to the combined effects of interfacial line tension and the midplane surface tension of anti-registered configurations (29,32,33), we do not independently measure either quantity here. Instead, Δ^∗^captures the experimentally observable outcome of their interplay. A closely related phenomenological approach to quantifying interleaflet coupling—based on comparing asymmetric and symmetric phase boundary locations along exchange-relevant composition trajectories—has been developed in parallel within our group using complementary lipid mixtures and experimental perturbations (Enoki and Heberle, submitted).

This framing also provides a natural connection to the concept of leaflet dominance introduced by Wang and London (25), with the sign of Δ^∗^ identifying the direction of dominance. Positive values of Δ^∗^ thus correspond to *separated leaflet dominance*, in which coupling to a phase-separating leaflet stabilizes demixing in the opposite leaflet, even though that leaflet would be uniformly mixed under symmetric conditions. Conversely, negative values of Δ^∗^correspond to *uniform leaflet dominance*, in which coupling to a uniformly mixed leaflet suppresses phase separation in a leaflet that would otherwise demix. Importantly, Δ^∗^extends this qualitative classification by providing a quantitative measure of the effective strength of dominance in either direction.

Although Δ^∗^ reflects the combined effects of both intraleaflet and interleaflet interactions, several observations are consistent with greater in-plane line tension in the DPPC/14:1-PC/Chol system. In particular, the larger value of *s*^∗^ for this mixture indicates a broader Ld+Lo coexistence region under symmetric conditions, consistent with an increased interfacial line tension associated with the larger expected thickness mismatch between coexisting phases, as shown previously (58,59). This interpretation is further supported by the observation that modulated phase morphologies were most frequent in DPPC/16:1-PC/Chol aGUVs and only rarely found in DPPC/14:1-PC/Chol (Fig. 3a). Such stripe-like or labyrinthine domain patterns are known to arise in regimes of low effective line tension, where the energetic cost of forming Ld/Lo interfaces is sufficiently small that macroscopic phase separation is replaced by patterned domain morphologies (60). The preferential appearance of modulated phases in the 16:1-PC system therefore provides independent evidence for lower effective line tension relative to the 14:1-PC system. This contrast further differentiates the two mixtures and is consistent with their distinct values of Δ^∗^.

At the same time, we cannot rule out contributions from changes in midplane surface tension, especially given our observation of three-phase coexistence in 14:1-PC aGUVs but not in the 16:1-PC system. Disentangling the relative roles of these factors will require additional experimental constraints, for example through systematic variation of lipid chain structure, possibly combined with direct measurements of midplane interfacial energetics (36). More broadly, by providing a direct, quantitative comparison between symmetric and asymmetric phase boundaries, the parameter Δ^∗^ offers a unifying framework for connecting phenomenological leaflet dominance concepts with coupled-leaflet theories, and for assessing the effective strength of interleaflet coupling in complex lipid mixtures.

### 4.2 Comparison with previous asymmetric bilayer studies

As discussed above, asymmetric phase boundaries emerge from the interplay of multiple energetic contributions, including intraleaflet interactions that govern in-plane Ld/Lo line tension and interleaflet interactions that contribute to the midplane surface tension of anti-registered configurations. Within this general framework, it is useful to ask how the phenomenology observed here compares to previous experimental studies of asymmetric bilayers. In particular, prior work reveals systematic trends in how lipid structure modulates the extent to which interleaflet coupling stabilizes or suppresses phase separation under asymmetric conditions.

Enoki and Feigenson reported that DSPC/18:1-PC/Chol aGUVs remained phase-separated even when essentially all of the outer leaflet DSPC was exchanged for 18:1-PC (corresponding to *ɛ*^∗^≈ 1), indicating strong stabilization of demixing by the opposing leaflet (26). In a separate study, the same authors showed that reducing the effective segregation strength—by replacing approximately half of the 18:1-PC with POPC—led to the emergence of an asymmetric phase boundary at *ɛ*^∗^ ≈ 0.8, which shifted steadily toward lower values with further replacement until macroscopic phase separation was completely abolished (27). These observations illustrate how changes in lipid structure can tune the extent to which interleaflet coupling stabilizes or suppresses phase separation under asymmetric conditions.

A consistent dependence on acyl chain length is also evident in the asymmetric LUV study of Wang and London, who examined the stability of ordered domains in outer leaflets composed of a saturated PC (with varying acyl chain length), 18:1-PC, and cholesterol—compositions that exhibit Ld+Lo coexistence in symmetric vesicles—opposite a uniformly mixed 18:1-PC/Chol leaflet (25). Across the temperature range examined, decreasing the saturated-chain length progressively destabilized outer leaflet domain formation relative to symmetric controls, with longer-chain lipids (DSPC and DPPC) retaining ordered domains over a broader temperature range than shorter-chain lipids (15:0-PC and DMPC). These results indicate that reduced chain-length mismatch weakens the ability of one leaflet to sustain phase separation in the presence of an opposing uniformly mixed leaflet, consistent with diminished interleaflet coupling strength. This trend qualitatively parallels the behavior observed here, in which decreasing acyl-chain mismatch correlates with increased susceptibility to suppression of macroscopic phase separation under asymmetric conditions at fixed temperature. Together, these studies support the view that lipid chain structure strongly modulates the effective strength of interleaflet coupling, as reflected in shifts of asymmetric phase boundaries relative to their symmetric counterparts. Within the present framework, such shifts are naturally captured by the parameter Δ^∗^, which reports how strongly interleaflet coupling biases leaflet miscibility without requiring attribution to a single microscopic interaction.

### 4.3 Mapping hemifusion pathways onto coupled-leaflet phase diagrams

To place our experimental observations within the framework of coupled-leaflet theories, it is useful to explicitly map the hemifusion-induced compositional perturbation onto theoretical leaflet-leaflet phase diagrams. To this end, we consider the illustrative phase diagrams shown in Fig. 6, which summarize key theoretical expectations in a form directly relevant to our hemifusion experiments. Fig. 6a-b depict both the free-energy landscape (upper triangular region) and corresponding leaflet-leaflet phase diagram (lower triangular region) as functions of the inner- and outer-leaflet order parameters. This compact representation exploits the symmetry of a flat bilayer, for which interchanging the inner and outer leaflet labels leaves the free energy unchanged, corresponding to reflection about the line *y* = *x*. In each diagram, selected tielines are drawn in the two-phase regions and the predicted three-phase region (i.e., R_D_+R_O_+AR_DO_) is shown as a shaded triangle. Blue arrows mark the approximate compositional path traversed during our hemifusion experiments, in which the outer leaflet order is progressively lowered while the composition of the inner leaflet remains fixed. Along this path, uppercase Roman numerals label the expected aGUV phenotypes, which are depicted schematically in Fig. 6c.

**Figure 6.**
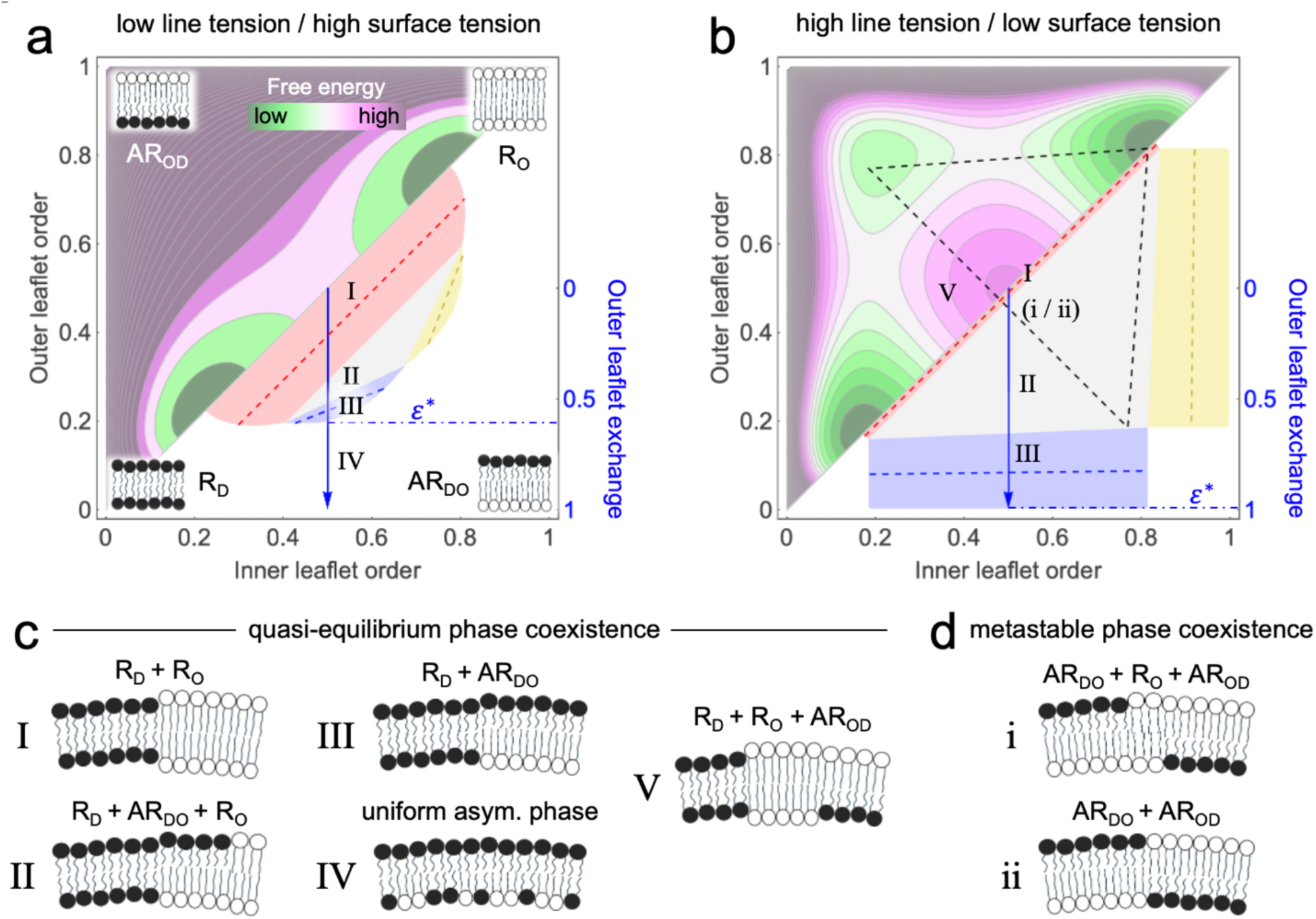
Coupled-leaflet free-energy landscapes and phase diagrams illustrating hemifusion trajectories and the emergence of anti-registered phases. Panels (a,b) show schematic free-energy landscapes (upper triangular regions) and the corresponding leaflet-leaflet phase diagrams (lower triangular regions) as functions of inner and outer leaflet order parameters; this compact representation exploits the symmetry of a flat bilayer, for which interchanging inner and outer leaflet labels leaves the free energy unchanged (reflection about the diagonal). Panel (a) illustrates a regime of relatively low line tension and high midplane surface tension, whereas panel (b) illustrates a regime of higher line tension and lower midplane surface tension, conditions that promote anti-registered (AR) configurations. Selected tielines are drawn in the two-phase regions (dashed lines), and the predicted three-phase coexistence region (R_D_ + R_O_ + AR_DO_) is indicated by a shaded triangle. Blue arrows denote the approximate compositional trajectory followed during hemifusion experiments, in which the outer leaflet order is progressively reduced while the inner leaflet composition remains fixed; the position of the asymmetric miscibility boundary, *ɛ*^∗^, is marked along this trajectory, illustrating its sensitivity to line tension and midplane surface tension. Roman numerals indicate expected aGUV phenotypes encountered along the trajectory. Panel (c) shows schematic representations of quasi-equilibrium coexistence phenotypes (I–V), including registered, mixed registered/anti-registered, and uniform asymmetric states, while panel (d) shows metastable AR coexistence phenotypes (i, ii) predicted to arise when the free-energy landscape contains stable AR minima.

The two leaflet-leaflet diagrams in Fig. 6 illustrate distinct regimes of Ld/Lo interfacial line tension and midplane surface tension, chosen to highlight how variations in these parameters alter both the free-energy landscape and the resulting asymmetric phase behavior. When the effective line tension is low and/or the midplane surface tension is high (Fig. 6a), the asymmetric miscibility boundary occurs at *ɛ*^∗^< 1, and a uniform asymmetric phase is accessible along the hemifusion pathway (phenotype IV). More generally, the location of *ɛ*^∗^ depends sensitively on the balance between interfacial line tension and midplane surface tension.

Increasing the effective line tension and/or decreasing the midplane surface tension stabilizes AR minima in the free-energy landscape, as illustrated in Fig. 6b. This stabilization has several consequences: (i) it expands the region of R_D_+R_O_+AR_DO_ coexistence; (ii) it shifts *ɛ*^∗^to larger values, potentially eliminating access to phenotype IV along the hemifusion trajectory; and (iii) it gives rise to additional regions of metastable coexistence involving AR phases, one of which is indicated by the dashed black triangle. Expected metastable phenotypes encountered along the hemifusion pathway are shown as lowercase Roman numerals in Fig. 6d. As discussed below, these schematic representations of distinct leaflet-coupled phase states provide a useful framework for interpreting the sporadic but reproducible appearance of AR phases in DPPC/14:1-PC/Chol aGUVs observed experimentally.

### 4.4 Anti-registered phases test the limits of coupled-leaflet theory

Our experiments reveal that anti-registered (AR) phases occasionally arise in DPPC/14:1-PC/Chol aGUVs generated by hemifusion. The most striking examples in our dataset are vesicles exhibiting coexisting AR_DO_+AR_OD_ phases, with or without an R_O_ domain, shown in Fig. 5b,c and illustrated schematically in Fig. 6d. Although AR phenotypes appeared in only a small minority of DPPC/14:1-PC/Chol vesicles (∼9%), they were highly reproducible, with at least one AR-containing vesicle observed in 7 of 9 preparations (Table S6). In contrast, no AR phases were detected in the DPPC/16:1-PC/Chol system. This compositional dependence is broadly consistent with the theoretical framework of Williamson and Olmsted (WO), which predicts that metastable AR coexistence emerges only when hydrophobic thickness mismatch is sufficiently large to produce true AR minima in the bilayer free-energy landscape, as in Fig. 6b (32,56,57).

One possible explanation for the appearance of metastable AR phases is that they originate before hemifusion, during the temperature quench that immediately follows electroformation of GUVs. In this scenario, some GUVs could become kinetically trapped in the AR basins rather than reaching the equilibrium R_D_+R_O_ coexistence. Such trapping is plausible because theoretical analyses predict that, immediately after a quench, AR instabilities can grow faster than registered ones when hydrophobic mismatch is sufficiently strong. However, this interpretation also presents a challenge: existing models generally predict that metastable AR domains should be replaced over time by registered domains through a nucleation-driven process (56). The observation of large, highly coarsened AR domains in our experiments therefore suggests that, if AR phases do arise during the temperature quench, they must be unusually long-lived, persisting well beyond the timescales typically expected for escape from AR metastability.

An alternative hypothesis is that the symmetric pre-hemifusion GUVs achieved equilibrium R_D_+R_O_ coexistence, and that hemifusion itself perturbed some of these vesicles into the regime of metastable AR coexistence. This scenario is more difficult to reconcile with existing theory, since it would require the hemifusion process to displace the system from a low-energy registered state into a higher-energy AR state. For this to occur, hemifusion would need to carry the bilayer through regions of composition space where the AR minima become kinetically accessible despite being thermodynamically disfavored. Whether such pathways are physically realistic is uncertain and not yet addressed by existing coupled-leaflet models, but the present observations provide constraints that may help refine theoretical models of how leaflet perturbations couple to lateral organization.

In this regard, one feature of our dataset is particularly striking: although WO theory predicts two metastable AR-AR-R coexistence regions—one associated with the R_O_ minimum and one with the R_D_ minimum—all nine vesicles exhibiting the AR-AR-R phenotype contained R_O_, and none contained R_D_. The reasons for this selective accessibility are not yet clear. A slight bias of the pre-hemifusion composition toward the R_O_ minimum could conceivably funnel vesicles into the AR_DO_+AR_OD_+R_O_ region. Alternatively, it is possible that coarsening from metastable AR-AR-R to the ultimate registered equilibrium state proceeds more rapidly when the registered phase is disordered, thereby making AR_DO_+AR_OD_+R_D_ more transient and therefore less likely to be observed. More generally, the absence of AR_DO_+AR_OD_+R_D_ vesicles may reflect the restricted compositional trajectory imposed by the hemifusion geometry or may indicate that additional physical factors—such as cholesterol redistribution, hemifusion-induced differential leaflet stress, or leaflet-specific mechanical constraints—play roles not yet incorporated into current coupled-leaflet theories and therefore represent important constraints for future theoretical development.

To summarize, although many aspects of our observations resonate with theoretical predictions, important discrepancies remain. In particular, the large size and longevity of coexisting AR domains suggest a more complex picture than represented in existing coupled-leaflet frameworks. Ultimately, we view these discrepancies as opportunities: the DPPC/14:1-PC/Chol mixture appears to offer an experimental system in which metastable AR coexistence—long predicted but rarely observed—can be directly visualized. These data may therefore provide an experimental foothold for future theoretical development, particularly in understanding the role of metastable AR basins, kinetic trapping pathways, and the influence of cholesterol and mechanical asymmetry during leaflet-specific perturbations. Similar perturbations occur in cells, where localized lipid insertion at membrane contact sites can transiently enrich a single leaflet before the bilayer fully equilibrates. Our hemifusion experiments provide a controlled analogue of this process, offering a useful model for exploring how such asymmetric inputs reshape lateral membrane organization.

### 4.5 Exploiting the highly variable asymmetry of hemifusion-generated aGUVs

Although the hemifusion technique is still too new to draw firm conclusions, a high degree of vesicle-to-vesicle variability in lipid asymmetry within the resulting population of aGUVs appears to be the rule. In addition to this study, we are aware of six published reports of aGUVs prepared by hemifusion where exchange was estimated from fluorescence intensity measurements (26,27,37,46-48). The aGUVs in these studies incorporate a variety of lipids (various PCs, PS, PIP(4,5)P_2_, cholesterol) and fluorophores (DiD, TF-PC, TF-PS, TF-PI(4,5)P_2_) and span a range of complexity (one to four lipid components), phase states (uniform Ld, Ld+Lo, Ld+Lβ), hemifusion initiation agents (Ca^2+^ and Mg^2+^), and solution conditions (low vs. high ionic strength). Together, these studies suggest that vesicle-to-vesicle variability in lipid exchange is an intrinsic feature of the hemifusion process, rather than a technical shortcoming.

Importantly, this intrinsic variability in lipid exchange is superimposed on a second, distinct contribution arising from uncertainty in fluorescence-based measurements of exchange. As discussed previously, the major source of error in the exchange measurement is variability in probe concentration in the initial symmetric GUVs (for the probe-exit experiment) and control GUVs used to calculate exchange (for both probe-exit and probe-entry experiments) (37). Comparing the fluorescence intensity of the symmetric control GUVs in the aforementioned studies, one interesting and useful observation that emerges is that some probes (TF-PC, TF-PS) seem to have a narrower intensity distribution than others (DiD, TF-PI(4,5)P_2_), suggesting that the former may yield more precise measurements of exchange. However, regardless of which probe is used, the intensity spread in aGUVs is almost always greater (and often much greater) than in the control GUVs; the lone exception is TF-PI(4,5)P_2_, which showed an unusually broad distribution of intensity in control GUVs that the authors attributed to increased fragility of PI(4,5)P_2_-containing vesicles (48). Although some of this additional spread may come from MLVs that escaped detection, as well as some vesicles undergoing full fusion rather than hemifusion, we believe that a substantial fraction of this additional spread can be attributed to variability in the amount of exchange. This is plausible given that differently sized vesicles may take shorter/longer periods of time to fully exchange their outer leaflets, and that initiation of exchange in individual GUVs likely occurs at different times and throughout the Ca^2+^ incubation period as GUVs continue to settle toward the SLB.

Some authors have emphasized the advantages of excluding vesicles that are not sufficiently asymmetric from subsequent analysis (26). The choice to focus on a subset of highly asymmetric GUVs is understandable from a biological standpoint, given the strong asymmetries (e.g., in lipid headgroups and chain unsaturation) known to exist in the PM (13). While we agree that highly asymmetric vesicles hold special significance, the large variability in outer leaflet exchange inherent to the hemifusion method can also be advantageous.

Focusing exclusively on fully exchanged vesicles is analogous to characterizing a protein solely by one favored conformation, rather than by the ensemble of states it can access. Indeed, it is unlikely that the highly asymmetric PM composition reported in Lorent et al. (13) represents the only physiologically relevant membrane state. Cells can partially or transiently relax membrane asymmetry through the activation of scramblases (61,62), suggesting that the degree of membrane asymmetry may itself be a dynamic variable under cellular control. Analyzing all asymmetric GUVs—including those with lesser amounts of exchange—therefore allows membrane properties (e.g., phase state, order, permeability, bending stiffness) to be mapped continuously as a function of asymmetry, providing direct insight into the nature of interleaflet coupling. In this light, the population of aGUVs produced by hemifusion can be viewed as sampling a trajectory through asymmetric composition space, revealing how properties such as macroscopic phase separation evolve as lipids in one leaflet are progressively replaced by a different set of lipids.

### 4.6 Interpreting and managing compositional uncertainty in hemifusion-generated aGUVs

Because hemifusion-generated aGUVs are increasingly used to probe leaflet-specific membrane properties, it is essential to understand how experimental design choices influence the uncertainty in inferred lipid asymmetry. In this work, we treat vesicle-to-vesicle variability in fluorescence intensity as the dominant source of uncertainty, arising from factors including instrumentation (e.g., spatial and temporal fluctuations in illumination intensity) (63), probe photostability, and intrinsic compositional variability even among nominally identical GUVs (64,65). Phase-separated vesicles are further subject to uncertainty associated with differences in the apparent phase fraction sampled in different confocal slices. These contributions are captured phenomenologically by the parameter *σ*_*F*_, which quantifies vesicle-to-vesicle variability in fluorescence intensity for a given probe and experimental configuration. Since aGUVs are prone to each of these sources of uncertainty in addition to others discussed below, vesicle-to-vesicle variation sets a lower bound on the precision with which outer-leaflet exchange can be inferred from Eqs. 1 and 2. The data in Tables S1–S3 therefore provide lower-bound estimates for *σ*_*F*_ of 10–25%.

A recent study from our group used an empirical maximum likelihood approach to estimate the compositional uncertainty of binary DPPC/DOPC aGUVs (37). In that case, symmetric GUVs that initially showed gel + fluid phase separation became homogeneously mixed when ≈ 60% of the outer leaflet DPPC was replaced with DOPC, resulting in two vesicle populations (i.e., phase-separated and uniform). As in the present study, some overlap in the distributions was observed, but the number of data points was too small to produce robust histograms for data fitting. Instead, the number of vesicles in the overlapping region was compared to simulated distributions to determine the most likely value of *σ*_*ɛ*_*obs*__, which was found to be 18% for the exiting probe and 12% for the entering probe. This analysis did not attempt to account for how the uncertainty might depend on the mode of probe exchange (i.e., exit vs. entry) or extent of exchange. It was also assumed that all values of exchange are equally probable, which is unlikely to generally be true, though experimental conditions can clearly be found such that this assumption is approximately met.

We developed a quantitative model to address the shortcomings mentioned above. Our starting point was an error propagation analysis of the equations used to calculate exchange fraction from GUV intensities (Supporting Information, Section S5.2). This analysis leads to three non-intuitive predictions, summarized in Fig. S8b. First, when exchange is inferred from the loss of a probe exiting the GUV, the associated uncertainty decreases with increasing exchange. Second, when exchange is inferred from a probe entering the GUV from the SLB, the uncertainty instead increases with increasing exchange. Third, if the fluorescence uncertainty *σ*_*F*_is independent of probe transfer direction, probe-entry experiments always yield more precise estimates of exchange than probe-exit experiments. The coupled-distributions model builds directly on the error-propagation analysis and provides a framework for assessing these predicted trends in experiment. The fact that the experimentally observed *ɛ*_*obs*_ distributions are well described by the model using a single framework for both probe-exit and probe-entry data lends support to our interpretation of the dominant sources of compositional uncertainty. It is also notable that a previous study using the same hemifusion approach reported a value of *σ*_*ɛobs*_ for the exiting probe that is 1.5-fold greater than that of the entering probe, in qualitative agreement with our prediction (37).

The magnitude of *σ*_*F*_ obtained from fitting the distributions (30–40%, Table 1) is larger than the lower-bound estimate based on the spread in fluorescence intensity seen in symmetric vesicles (10–25%, Tables S1-S3). As mentioned above, a fundamental difference between symmetric and asymmetric vesicles is that hemifusion-generated aGUVs exhibit substantially greater intrinsic compositional variability than conventionally prepared symmetric GUVs. Because probe intensity is often strongly dependent on composition due to fluorophore environmental sensitivity (66,67), the large compositional variability of aGUVs likely contributes to greater variability in probe intensity. Another confounding factor is the possibility of full fusion, which was previously observed in a small subset of hemifusion-generated aGUVs under similar calcium concentrations (26). Values of *ɛ*_*obs*_greater than 1 can arise when both GUV leaflets are available for exchange with the SLB. By contrast, the presence of multilamellar vesicles would have the opposite effect, resulting in negative apparent exchange in probe-exit measurements. We also note that the fitted values of *σ*_*F*_ differ for the probe-exit and probe-entry experiments in the DPPC/16:1-PC/Chol system (Table 1). Additional data will be required before drawing firm conclusions about how *σ*_*F*_might depend on the direction of probe transfer.

The values for *σ*_*ɛobs*_ obtained from our analysis are substantially larger than values derived from maximum likelihood estimation as discussed previously (37). An additional factor contributing to uncertainty in the present study is our use of a single fluorescent probe to quantify *ɛ*_*obs*_. In several previous hemifusion studies, spectrally distinct fluorophores were incorporated into the pre-hemifusion GUVs and SLB, enabling simultaneous and independent estimates of exchange from probe exit and probe entry (26,27,37,46). In our experiments, a second (red) probe was included solely for visualization of phase behavior and was deliberately kept at low concentration to minimize photophysical artifacts—such as energy transfer—that could compromise quantitative intensity-based measurements. As a result, *ɛ*_*obs*_was determined from a single probe (TFPC) in each experiment. Typically, the average of two independent measurements of comparable error will reduce the uncertainty by a factor of 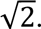 However, as described above, propagation of uncertainty reveals that the error associated with probe entry is generally smaller than that of probe exit when *σ*_*F*_ is similar for both modes of probe transfer (Section S5 and Fig. S8b). In this case, the uncertainty associated with the average of the two measurements will be *greater* than that of the probe-entry measurement alone until the exchange fraction reaches a value of ∼ 0.73. It is therefore unlikely that the smaller uncertainties reported in previous hemifusion studies arise from averaging probe-exit and probe-entry measurements. Rather, they likely reflect the use of selection criteria, in which aGUVs were excluded from subsequent analysis if the value of *ɛ*_*obs*_measured from exiting and entering probes disagreed by more than a specified threshold (e.g., 20%), or if either *ɛ*_*obs*_ value was deemed unphysical (i.e., < 0 or > 100% exchange) (26,27,37,46). Applying these criteria may have the effect of eliminating MLVs or fully fused vesicles that are otherwise difficult to identify. We chose to include all vesicles in our analysis with the goal of determining the true uncertainty of exchange measurements. This information, and the methodology for determining error introduced here, should prove valuable in the continued development and optimization of the hemifusion technique.

### 4.7 Limitations arising from unknown cholesterol distribution

While we quantify hemifusion-facilitated *phospholipid* exchange indirectly from the intensity of fluorescent lipid probes, we do not directly measure the final cholesterol concentration in aGUVs, nor do we know its distribution between the two leaflets. Given its rapid transbilayer flip-flop (sub-microsecond to millisecond timescales), cholesterol is expected to rapidly equilibrate its chemical potential between the inner and outer leaflets, in contrast to phospholipids, which remain essentially kinetically trapped in their respective leaflets on the timescale of our experiments. Consistent with this view, recent thermodynamic models of asymmetric bilayers have been formulated under the assumption of equal cholesterol chemical potential in the two leaflets (68).

Whether hemifusion results in a net transfer of cholesterol into or out of the GUV depends on the relative chemical potentials of cholesterol in the SLB and the pre-hemifusion GUV. In principle, this complication could be minimized by matching cholesterol chemical potentials rather than concentrations, although such data are not currently available for the lipid mixtures studied here. Measurements of cholesterol chemical potential in the related DPPC/DOPC/Chol system (69), together with the compositions of our SLBs and GUVs, suggest that a net flux of cholesterol from the SLB into the GUV is plausible under our experimental conditions. Direct determination of cholesterol content in individual vesicles, however, remains technically challenging. Additional uncertainty may arise if differential leaflet stresses accumulate during hemifusion, as such stresses can alter cholesterol chemical potential and bias its interleaflet distribution (70). Each of these factors could, in principle, contribute to the changes in phase behavior observed in highly asymmetric aGUVs.

Despite these limitations, experiments of the type presented here provide essential constraints for thermodynamic models of interleaflet coupling. By quantifying how phase behavior evolves as lipid asymmetry is systematically tuned, even in the absence of direct measurements of all molecular degrees of freedom, such data offer a critical bridge between theoretical predictions and experimentally accessible observables in asymmetric membranes.

## Author contributions

F.A.H. conceived of the project and obtained funding. K.B.K-C. and F.A.H. designed the research. K.B.K-C. and A.M.C. performed the experiments. K.B.K-C. and F.A.H. analyzed the data and wrote the manuscript.

## Acknowledgements

We thank Thais Enoki and Haden Scott for help with the hemifusion protocol. We thank John Williamson, Peter Olmsted, and Thais Enoki for helpful discussions. This research was supported by NIH grant R01 GM138887 (to F.A.H). K.B.K-C. was supported by the Graduate Advancement & Training Education (GATE) fellowship from the University of Tennessee-Oak Ridge Innovation Institute (UT-ORII). Assistance from ChatGPT-5.2 (Open AI, https://chat.openai.com/) was used to improve the clarity, grammar, and conciseness of the manuscript text.

