## Supporting Information for "Quantifying interleaflet coupling of phase behavior and observing anti-registered phases in asymmetric lipid bilayers"

### Table of Contents:

S1. Overview
S2. Validation of symmetric GUV controls
    Figures S1-S4
    Tables S1-S3
S3. Asymmetric GUVs: DPPC/16:1-PC/Chol system
    Figures S5-S6
    Tables S4-S5
S4. Asymmetric GUVs: DPPC/14:1-PC/Chol system
    Figure S7
    Table S6
S5. Extended modeling and uncertainty analysis
    Text S5.1 (coupled-distributions model)
    Text S5.2 (error propagation derivation, Eqs. S1-S8)
    Figures S8-S9
S6. Supplementary control studies
    Figure S10
    Tables S7-S8

**Figure and Table List**

**Section S2. Symmetric GUV controls**

- **Figure S1.** TFPC intensity and phase fraction data for symmetric DPPC/16:1-PC/Chol GUVs used for probe-exit calculations.
- **Figure S2.** Correlations of vesicle diameter with intensity and Ld phase fraction for symmetric DPPC/16:1-PC/Chol GUVs.
- **Figure S3.** TFPC fluorescence intensity distributions for symmetric DPPC/16:1-PC/Chol GUVs used for probe-entry calculations.
- **Figure S4.** TFPC intensity and phase fraction data for symmetric DPPC/14:1-PC/Chol GUVs used for probe-entry calculations.
- **Table S1.** Summary of symmetric DPPC/16:1-PC/Chol GUVs (probe-exit mode).
- **Table S2.** Summary of symmetric DPPC/16:1-PC/Chol GUVs (probe-entry mode).
- **Table S3.** Summary of symmetric DPPC/14:1-PC/Chol GUVs (probe-entry mode).

**Section S3. Asymmetric GUVs: DPPC/16:1-PC/Chol**

- **Figure S5.** Ld phase fractions of individual phase-separated aGUVs across preparations for probe-exit and entry modes.
- **Figure S6.** Modulated phase morphologies in DPPC/16:1-PC/Chol aGUVs.
- **Table S4.** Summary of 16:1-PC aGUVs (probe-exit mode).
- **Table S5.** Summary of 16:1-PC aGUVs (probe-entry mode).

**Section S4. Asymmetric GUVs: DPPC/14:1-PC/Chol**

- **Figure S7.** Ld phase fractions of two-phase aGUVs by preparation.
- **Table S6.** Summary of 14:1-PC aGUVs (probe-entry mode).

**Section S5. Extended modeling and uncertainty analysis**

- **Text S5.1.** Coupled-distributions model.
- **Figure S8.** Coupled-distributions model and uncertainty propagation
- **Text S5.2.** Derivation of uncertainty in calculated exchange fraction (Eqs. S1–S8).
- **Figure S9.** Sensitivity of aGUV distribution model to uncertainty in the parameter λ.

**Section S6. Supplementary control studies**

- **Figure S10.** Confocal *z*-stack images confirming that equatorial Ld phase fractions represent overall vesicle phase fractions.
- **Table S7.** Symmetric DPPC/14:1-PC/Chol GUVs for locating ternary phase boundary.
- **Table S8.** Symmetric DPPC/16:1-PC/Chol GUVs for locating ternary phase boundary.

### S1. Overview

This Supporting Information provides additional data and analyses that complement the main text. **Sections S2–S4** present the complete datasets for symmetric and asymmetric giant unilamellar vesicles (GUVs) used in this study, including fluorescence intensity distributions and phase fraction statistics for both DPPC/16:1-PC/Chol and DPPC/14:1-PC/Chol mixtures. **Section S5** details the derivation and application of the coupled-distributions model used to determine the asymmetric phase boundary location and to quantify compositional uncertainty in individual aGUVs. **Section S6** provides supplementary control studies including (i) *z*-stack imaging to verify equatorial phase fraction accuracy and (ii) data from symmetric GUVs for locating ternary phase boundary locations. Together, these materials document data reproducibility, describe model implementation, and supply methodological transparency for all reported results.

### S2. Symmetric GUV controls.

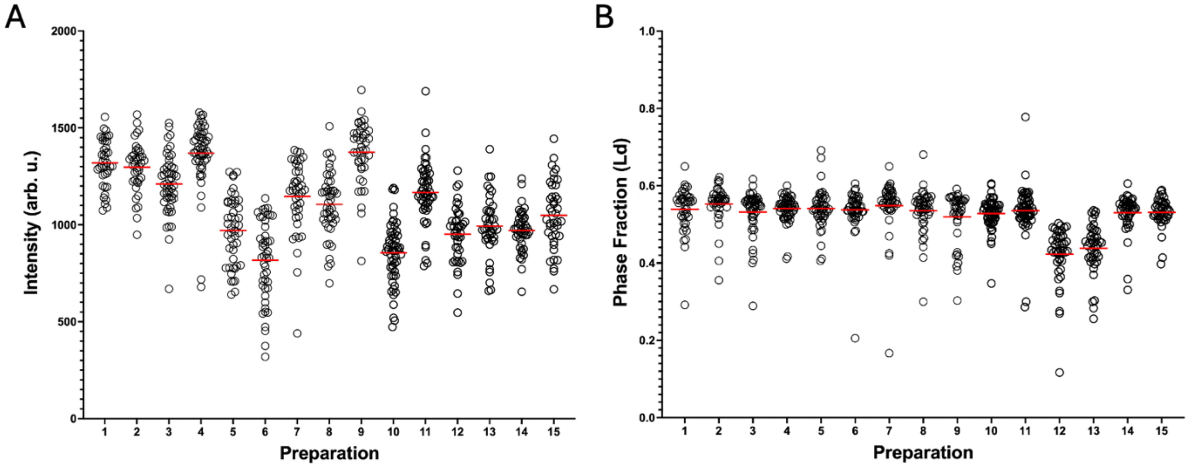


**Figure S1.** Distributions of TFPC intensity (A) and Ld phase area fraction (B) for symmetric DPPC/16:1-PC/Chol GUVs (used for quantifying exchange in probe-exit mode) across preparations. Horizontal red lines are means. All vesicles were phase separated.

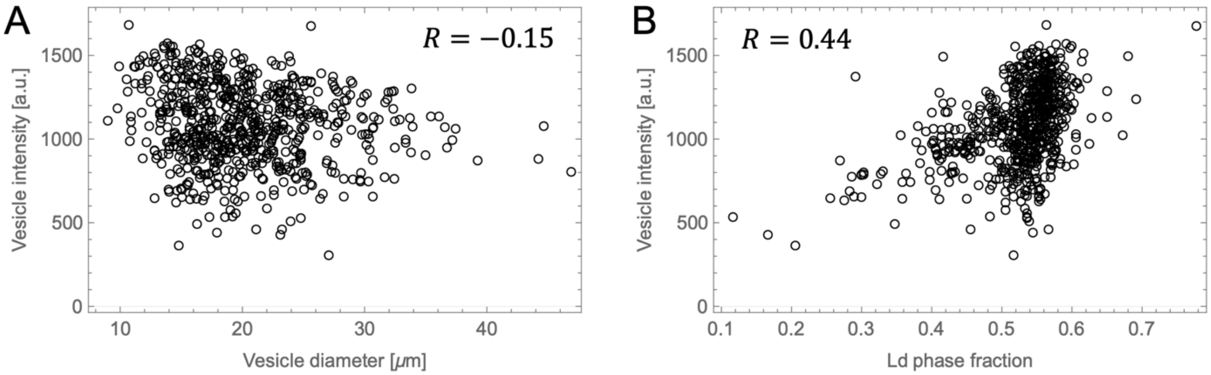


**Figure S2.** Correlations between average TFPC intensity and (A) vesicle diameter, or (B) Ld-phase fraction (B) for symmetric DPPC/16:1-PC/Chol GUVs.

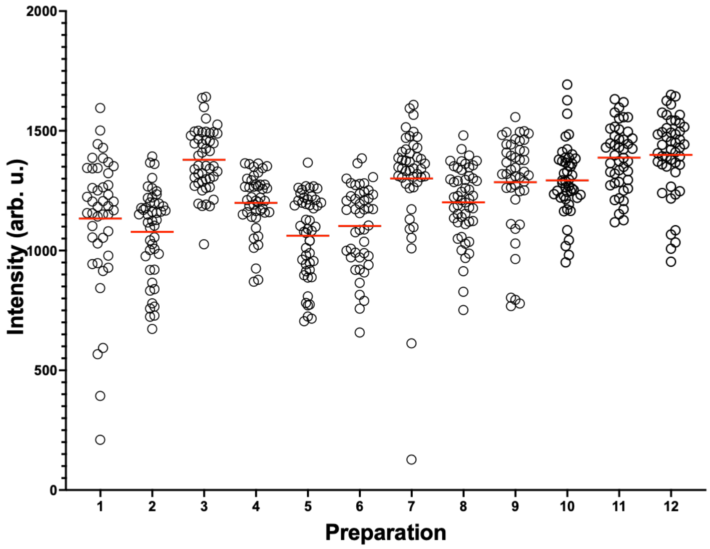


**Figure S3.** TFPC intensity distributions for symmetric DPPC/16:1-PC/Chol GUVs used for probe-entry calculations in asymmetry experiments. Horizontal red lines are means. All vesicles were phase-separated.

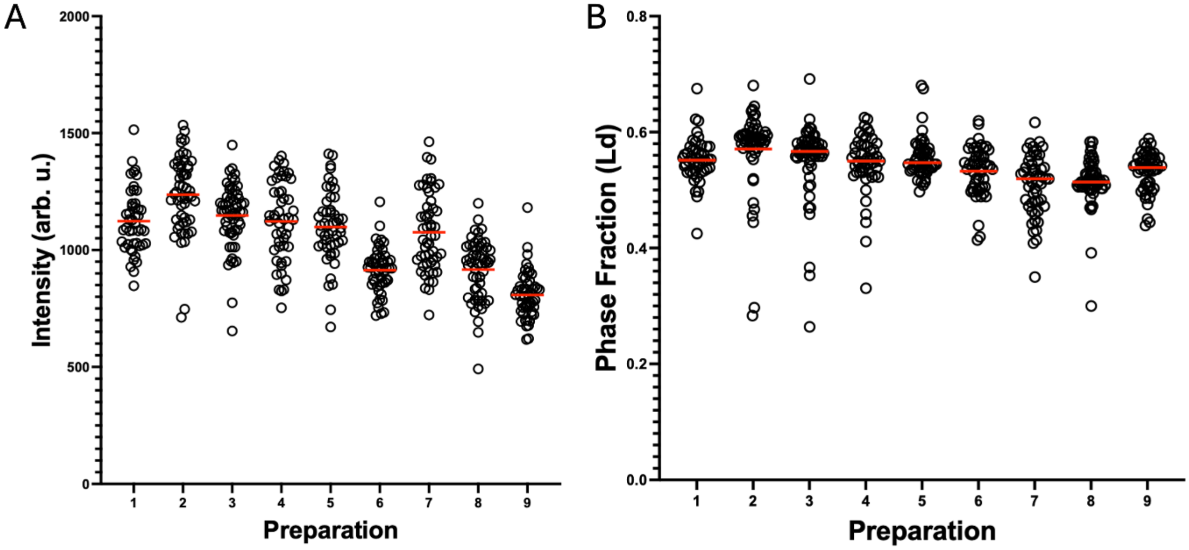


**Figure S4.** TFPC intensity (A) and Ld phase area fraction (B) for symmetric DPPC/14:1-PC/Chol GUVs confirming reproducible Lo + Ld coexistence. Horizontal red lines are means. All vesicles were phase separated.

**Table S1.** Summary of symmetric DPPC/16:1-PC/Chol GUVs used in probe-exit experiments: number of vesicles analyzed (*N*), mean fluorescence intensity ($\bar{F}$) and relative uncertainty ($\sigma_{\bar{F}}/\bar{F}$) of TFPC, mean diameter ($d$), average Ld phase area fraction ($f_{Ld}$), and relative uncertainty in Ld area fraction ($\sigma_{f_{Ld}}/f_{Ld}$).

| Prep | $N$ | $\bar{F}$ | $\sigma_{\bar{F}}/\bar{F}$ (%) | $d$ ($\mu m$) | $f_{Ld}$ | $\sigma_{f_{Ld}}/f_{Ld}$ (%) |
|---|---|---|---|---|---|---|
| DOPC | 30 | 1040 ± 106 | 10.2 | 20 ± 6 | -- | -- |
| 1 | 40 | 1320 ± 122 | 9.2 | 19 ± 4.4 | 0.54 ± 0.059 | 11.0 |
| 2 | 40 | 1300 ± 130 | 10.0 | 27 ± 6.1 | 0.55 ± 0.051 | 9.2 |
| 3 | 50 | 1210 ± 155 | 12.8 | 23 ± 4.5 | 0.53 ± 0.057 | 10.8 |
| 4 | 53 | 1370 ± 168 | 12.3 | 19 ± 4.2 | 0.54 ± 0.033 | 6.2 |
| 5 | 47 | 971 ± 174 | 18.0 | 21 ± 4.8 | 0.54 ± 0.051 | 9.4 |
| 6 | 47 | 817 ± 209 | 25.6 | 19 ± 4.8 | 0.54 ± 0.056 | 10.4 |
| 7 | 42 | 1150 ± 188 | 16.4 | 20 ± 5 | 0.55 ± 0.073 | 13.4 |
| 8 | 47 | 1110 ± 165 | 14.8 | 18 ± 5.1 | 0.54 ± 0.056 | 10.4 |
| 9 | 42 | 1370 ± 161 | 11.7 | 16 ± 3.6 | 0.52 ± 0.067 | 12.9 |
| 10 | 57 | 852 ± 158 | 18.5 | 19 ± 4.5 | 0.53 ± 0.039 | 7.4 |
| 11 | 53 | 1170 ± 159 | 13.6 | 26 ± 3.5 | 0.54 ± 0.065 | 12.1 |
| 12 | 43 | 952 ± 146 | 15.3 | 20 ± 4 | 0.42 ± 0.074 | 17.4 |
| 13 | 42 | 993 ± 154 | 15.5 | 18 ± 4.0 | 0.44 ± 0.064 | 14.7 |
| 14 | 42 | 971 ± 108 | 11.1 | 19 ± 4.0 | 0.53 ± 0.050 | 9.4 |
| 15 | 41 | 1050 ± 174 | 16.5 | 19 ± 5.0 | 0.53 ± 0.038 | 7.19 |
| Mean | 46 | 1110 ± 182 | 14.8 ± 4.1 | 20 ± 3 | 0.52 ± 0.039 | 10.8 ± 3.0 |

**Table S2.** Summary of symmetric DPPC/16:1-PC/Chol GUVs used as normalization controls for probe-entry calculations: number of vesicles analyzed (*N*), mean fluorescence intensity ($\bar{F}$) and relative uncertainty ($\sigma_{\bar{F}}/\bar{F}$) of TFPC, mean diameter ($d$), average Ld phase area fraction ($f_{Ld}$), and relative uncertainty in Ld area fraction ($\sigma_{f_{Ld}}/f_{Ld}$).

| Prep | $N$ | $\bar{F}$ | $\sigma_{\bar{F}}/\bar{F}$ (%) | $d$ ($\mu m$) | $f_{Ld}$ | $\sigma_{f_{Ld}}/f_{Ld}$ (%) |
|---|---|---|---|---|---|---|
| 1 | 44 | 1130 ± 279 | 24.6 | 19 ± 7.6 | 0.50 ± 0.12 | 23.1 |
| 2 | 46 | 1080 ± 186 | 17.2 | 19 ± 3.5 | 0.48 ± 0.10 | 21.1 |
| 3 | 47 | 1380 ± 131 | 9.5 | 24 ± 5.7 | 0.54 ± 0.033 | 6.2 |
| 4 | 43 | 1200 ± 120 | 10.4 | 22 ± 7.5 | 0.52 ± 0.027 | 5.2 |
| 5 | 49 | 1060 ± 175 | 16.5 | 19 ± 4.5 | 0.54 ± 0.053 | 9.8 |
| 6 | 45 | 1100 ± 170 | 15.7 | 19 ± 3.7 | 0.54 ± 0.035 | 6.5 |
| 7 | 44 | 1300 ± 250 | 19.2 | 22 ± 6.1 | 0.56 ± 0.039 | 7.1 |
| 8 | 50 | 1200 ± 150 | 12.9 | 21 ± 6.4 | 0.58 ± 0.052 | 9.0 |
| 9 | 44 | 1290 ± 205 | 16.0 | 21 ± 4.7 | 0.54 ± 0.070 | 12.9 |
| 10 | 46 | 1300 ± 150 | 11.6 | 22 ± 4.5 | 0.55 ± 0.086 | 8.6 |
| 11 | 45 | 1390 ± 135 | 9.8 | 21 ± 6.2 | 0.54 ± 0.032 | 6.0 |
| 12 | 47 | 1400 ± 170 | 12.0 | 21 ± 8.9 | 0.55 ± 0.045 | 8.2 |
| Mean | 46 | 1240 ± 124 | 14.6 ± 4.5 | 21 ± 1.6 | 0.54±0.026 | 10.3 ± 5.9 |

**Table S3.** Summary of symmetric DPPC/14:1-PC/Chol GUVs used as normalization controls for probe-entry calculations: number of vesicles analyzed (*N*), mean fluorescence intensity ($\bar{F}$) and relative uncertainty ($\sigma_{\bar{F}}/\bar{F}$) of TFPC, mean diameter ($d$), average Ld phase area fraction ($f_{Ld}$), and relative uncertainty in Ld area fraction ($\sigma_{f_{Ld}}/f_{Ld}$).

| Prep | $N$ | $\bar{F}$ | $\sigma_{\bar{F}}/\bar{F}$ (%) | $d$ ($\mu m$) | $f_{Ld}$ | $\sigma_{f_{Ld}}/f_{Ld}$ (%) |
|---|---|---|---|---|---|---|
| 1 | 46 | 1120 ± 138 | 12.3 | 17 ± 3.9 | 0.55 ± 0.039 | 6.9 |
| 2 | 53 | 1240 ± 167 | 13.5 | 18 ± 4.2 | 0.57 ± 0.071 | 12.5 |
| 3 | 59 | 1150 ± 136 | 11.9 | 23 ± 5.8 | 0.55 ± 0.065 | 11.7 |
| 4 | 50 | 1120 ± 169 | 15.1 | 24 ± 5.5 | 0.55 ± 0.052 | 9.5 |
| 5 | 50 | 1100 ± 150 | 14.1 | 21 ± 4.2 | 0.56 ± 0.035 | 6.4 |
| 6 | 50 | 914 ± 97.1 | 10.6 | 19 ± 3.3 | 0.53 ± 0.042 | 8.0 |
| 7 | 53 | 1080 ± 127 | 15.5 | 14 ± 2.8 | 0.51 ± 0.053 | 10.3 |
| 8 | 55 | 917 ± 132 | 14.4 | 16 ± 3.2 | 0.51 ± 0.043 | 8.4 |
| 9 | 51 | 807 ± 101 | 12.5 | 18 ± 3.1 | 0.53 ± 0.035 | 6.5 |
| Mean | 52 | 1050 ± 139 | 13.2 ± 1.6 | 19 ± 3.3 | 0.54 ± 0.021 | 8.9 ± 2.2 |

### S3. Asymmetric GUVs: DPPC/16:1-PC/Chol system

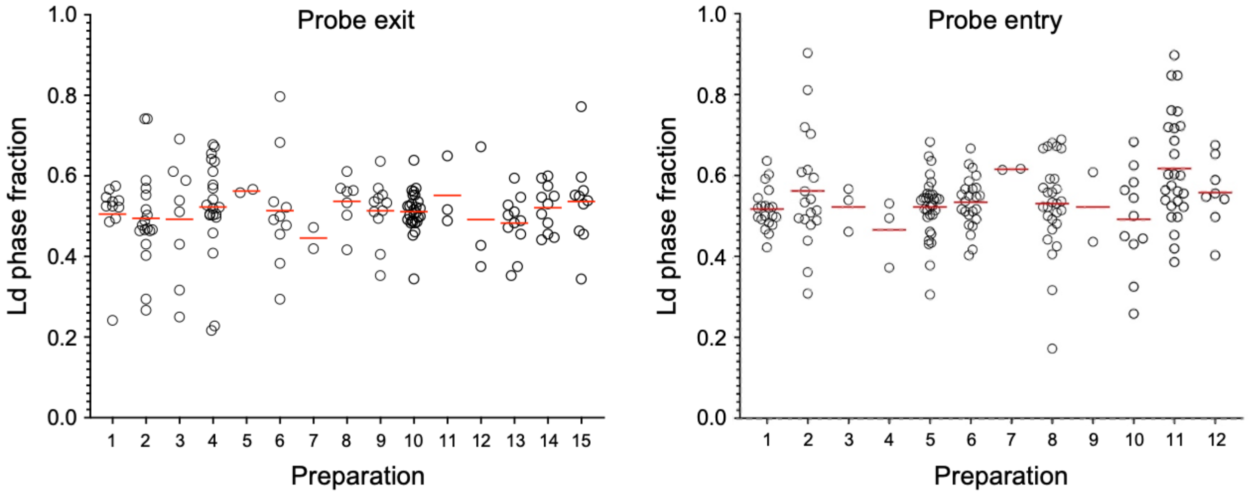


**Figure S5.** Ld phase fractions of individual phase-separated DPPC/16:1-PC/Chol aGUVs from probe-exit (left) and probe-entry (right) experiments plotted by preparation, demonstrating consistent phase fractions among replicates. Horizontal red lines are means.

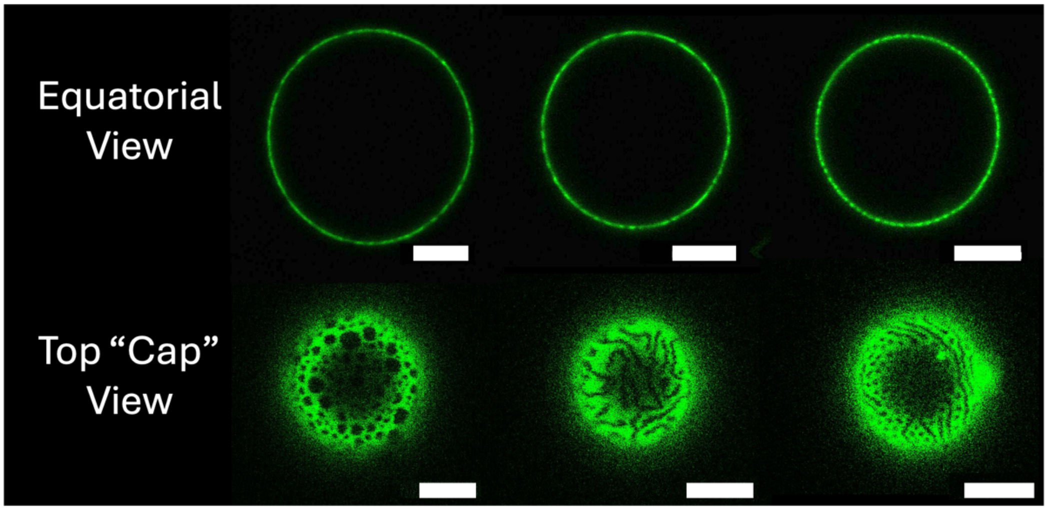


**Figure S6.** Examples of modulated phase morphologies in DPPC/16:1-PC/Chol aGUVs. Shown are three representative aGUVs exhibiting small and rounded or stripe-like domain patterns characteristic of modulated phases. For each vesicle, an equatorial confocal slice (top row) and a view of the vesicle cap (bottom row) are shown. These morphologies were observed across a wide range of outer leaflet exchange fractions. Scale bare are 5 μm.

**Table S4.** Data for DPPC/16:1-PC/Chol aGUVs obtained from probe-exit experiments: number of vesicles ($N$), number of phase-separated vesicles including modulated ($N_{2\phi}$), number of modulated vesicles ($N_{mod}$), Ld phase area fraction ($f_{Ld}$), relative uncertainty in Ld area fraction ($\sigma_{f_{Ld}}/f_{Ld}$).

| Prep | $N$ | $N_{2\phi}$ | $N_{mod}$ | $f_{Ld}$ | $\sigma_{f_{Ld}}/f_{Ld}$ (%) |
|---|---|---|---|---|---|
| 1 | 20 | 11 | 0 | 0.51 ± 0.091 | 18.1 |
| 2 | 32 | 18 | 0 | 0.49 ± 0.12 | 24.5 |
| 3 | 13 | 9 | 1 | 0.49 ± 0.15 | 30.6 |
| 4 | 26 | 24 | 3 | 0.52 ± 0.12 | 23.6 |
| 5 | 4 | 4 | 2 | 0.50 ± 0.10 | 20.6 |
| 6 | 14 | 10 | 0 | 0.51 ± 0.14 | 27.5 |
| 7 | 2 | 2 | 0 | 0.45 ± 0.037 | 8.4 |
| 8 | 8 | 7 | 0 | 0.54 ± 0.063 | 11.7 |
| 9 | 21 | 14 | 3 | 0.51 ± 0.077 | 15.0 |
| 10 | 34 | 30 | 3 | 0.51 ± 0.051 | 10.0 |
| 11 | 7 | 5 | 2 | 0.55 ± 0.086 | 15.1 |
| 12 | 6 | 3 | 0 | 0.49 ± 0.16 | 32.3 |
| 13 | 15 | 11 | 0 | 0.48 ± 0.070 | 14.5 |
| 14 | 16 | 13 | 2 | 0.52 ± 0.058 | 11.2 |
| 15 | 11 | 11 | 1 | 0.54 ± 0.11 | 20.5 |
| Mean | | | | 0.51 ± 0.026 | 18.9 ± 7.6 |

**Table S5.** Data for DPPC/16:1-PC/Chol aGUVs obtained from probe-entry experiments: number of vesicles ($N$), number of phase-separated vesicles including modulated ($N_{2\phi}$), number of modulated vesicles ($N_{mod}$), Ld phase area fraction ($f_{Ld}$), relative uncertainty in Ld area fraction ($\sigma_{f_{Ld}}/f_{Ld}$).

| Prep | $N$ | $N_{2\phi}$ | $N_{mod}$ | $f_{Ld}$ | $\sigma_{f_{Ld}}/f_{Ld}$ (%) |
|---|---|---|---|---|---|
| 1 | 24 | 22 | 2 | 0.52 ± 0.051 | 9.9 |
| 2 | 26 | 20 | 1 | 0.56 ± 0.14 | 25.1 |
| 3 | 4 | 3 | 0 | 0.52 ± 0.055 | 10.5 |
| 4 | 5 | 3 | 0 | 0.47 ± 0.083 | 17.8 |
| 5 | 50 | 39 | 7 | 0.52 ± 0.075 | 14.3 |
| 6 | 29 | 27 | 3 | 0.53 ± 0.063 | 11.7 |
| 7 | 3 | 2 | 0 | 0.62 ± 0.0020 | 0.3 |
| 8 | 34 | 29 | 1 | 0.53 ± 0.11 | 21.2 |
| 9 | 3 | 2 | 0 | 0.52 ± 0.12 | 23.3 |
| 10 | 16 | 11 | 0 | 0.49 ± 0.13 | 25.8 |
| 11 | 36 | 26 | 0 | 0.62 ± 0.13 | 21.7 |
| 12 | 17 | 9 | 1 | 0.56 ± 0.086 | 15.4 |
| Mean | | | | 0.54 ± 0.045 | 16.4 ± 7.6 |

### S4. Asymmetric GUVs: DPPC/14:1-PC/Chol system

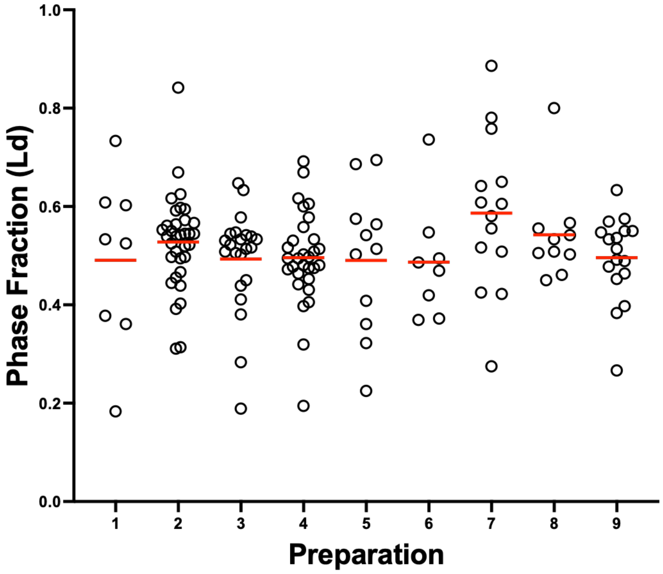


**Figure S7.** Ld phase fractions for aGUVs from DPPC/14:1-PC/Chol plotted by preparation. Horizontal red lines are means.

**Table S6.** Data for DPPC/14:1-PC/Chol aGUVs obtained from probe-entry experiments: number of vesicles ($N$), number of 2-phase vesicles including modulated ($N_{2\phi}$), number of 3-phase vesicles ($N_{3\phi}$), number of modulated vesicles ($N_{mod}$), Ld phase area fraction ($f_{Ld}$), relative uncertainty in Ld area fraction ($\sigma_{f_{Ld}}/f_{Ld}$).

| Prep | $N$ | $N_{2\phi}$ | $N_{3\phi}$ | $N_{mod}$ | $f_{Ld}$ | $\sigma_{f_{Ld}}/f_{Ld}$ (%) |
|---|---|---|---|---|---|---|
| 1 | 10 | 10 | 0 | 2 | 0.49 ± 0.17 | 35.5 |
| 2 | 41 | 34 | 2 | 0 | 0.53 ± 0.097 | 18.4 |
| 3 | 24 | 22 | 1 | 0 | 0.49 ± 0.10 | 21.1 |
| 4 | 35 | 31 | 1 | 1 | 0.49 ± 0.097 | 19.5 |
| 5 | 16 | 11 | 5 | 0 | 0.49 ± 0.15 | 30.1 |
| 6 | 12 | 9 | 3 | 1 | 0.49 ± 0.12 | 24.3 |
| 7 | 18 | 14 | 2 | 0 | 0.59 ± 0.16 | 27.0 |
| 8 | 10 | 10 | 0 | 0 | 0.54 ± 0.098 | 18.1 |
| 9 | 21 | 16 | 2 | 0 | 0.51 ± 0.066 | 12.9 |
| Mean | | | | | 0.51 ± 0.035 | 23.0 ± 7.0 |

### S5. Extended modeling and uncertainty analysis

#### S5.1 Coupled-distributions model

The substantial overlap in the distributions of uniform and phase-separated aGUVs (as in Fig. 3a) makes it difficult to determine the location of the asymmetric miscibility boundary ($\varepsilon^*$) by simple visual inspection. We therefore developed a coupled-distributions model to fit the experimental exchange data, described in Methods and expanded upon here. The salient features of the model are demonstrated in Fig. S8 and summarized here:

1. The probability $f(\varepsilon)$ for an aGUV to have a *true* exchange fraction $\varepsilon$ is assumed to be a function of a single adjustable parameter, $\lambda$, according to Eq. 3. Rather than representing a fixed physical property of the system, $\lambda$ parameterizes the stage of an evolving exchange process across the vesicle population. When $\lambda = 0$, all values of $\varepsilon$ are equally likely. Positive (negative) values of $\lambda$ shift the distribution toward smaller (larger) values of $\varepsilon$, respectively (Fig. S8a).

2. The *observed* distribution of exchange fractions arises from convolution of the true exchange distribution, $f(\varepsilon)$, with uncertainty in estimating exchange from confocal fluorescence measurements. We represent this uncertainty by a normal distribution, $g(\varepsilon_{obs}; \varepsilon, \sigma_F)$, which describes the probability of measuring an apparent exchange fraction $\varepsilon_{obs}$ for an aGUV with true exchange fraction $\varepsilon$ in probe-exit (Eq. 4a) or probe-entry (Eq. 4b) experiments. As shown in Section S5.2 below, error propagation analysis of Eqs. 1 and 2 reveals that the widths of these distributions, $\sigma_{\varepsilon_{obs}}$, depend on both $\varepsilon$ and the relative error associated with measuring aGUV fluorescence intensity, $\sigma_F$, as shown in Fig. S8b. In probe-exit experiments, the uncertainty $\sigma_{\varepsilon_{obs}}$ is largest at low $\varepsilon$ and decreases linearly with increasing $\varepsilon$; in contrast, in probe-entry experiments the uncertainty is smallest at low $\varepsilon$ and increases linearly with $\varepsilon$.

3. When the asymmetric composition space contains a phase boundary $\varepsilon^*$, the probability distributions of $\varepsilon_{obs}$ for uniform and phase-separated vesicles ($P^{1\phi}$ and $P^{2\phi}$, respectively) are given by Eqs. 5. These distributions are uniquely determined by the shared parameters $\lambda$, $\sigma_F$, and $\varepsilon^*$, and are therefore intrinsically coupled through both the underlying exchange distribution and the location of the asymmetric miscibility boundary. Fig. S8c,d demonstrates how $P^{1\phi}$ and $P^{2\phi}$ spread out with increasing $\sigma_F$ for the probe-exit (Fig. S8c) and probe-entry (Fig. S8d) experiments.

Importantly, the coupled-distributions framework enables all observed vesicles—including those with $\varepsilon_{obs}$ values outside the physical range [0,1]—to contribute information when fitting the model to experimental data. In practice, the model is fit simultaneously to the observed $\varepsilon_{obs}$ distributions of uniform and phase-separated vesicles, yielding estimates of $\varepsilon^*$, $\sigma_F$, and $\lambda$ that are consistent across both phenotypes and both probe-exchange modes. As demonstrated in the main text, this population-level approach provides a robust means of extracting asymmetric miscibility boundaries despite substantial uncertainty in $\varepsilon_{obs}$ at the level of individual aGUVs.

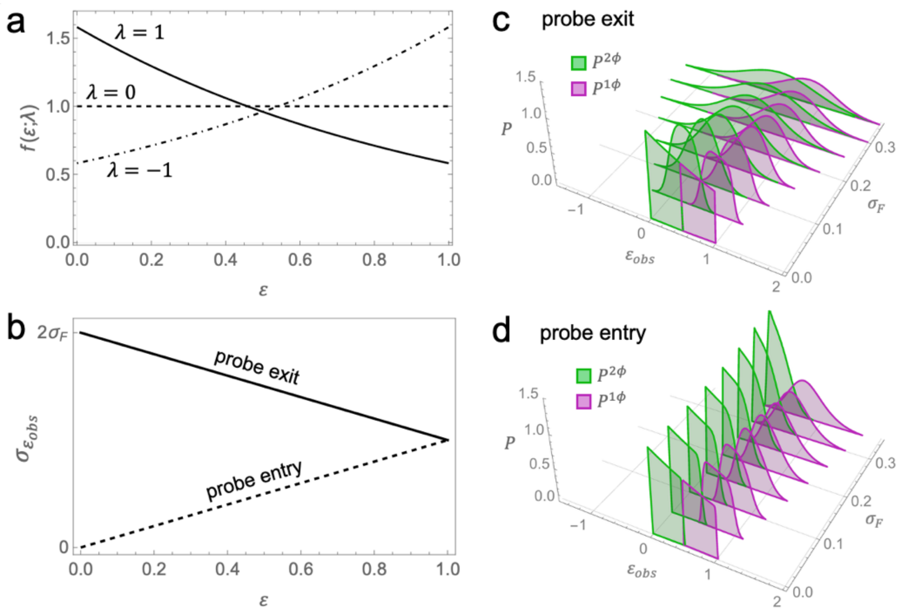


**Figure S8.** Coupled-distributions model and uncertainty propagation. (a) Model for the probability distribution of true exchange fraction, $f(\varepsilon)$ (Eq. 3), shown for three representative values of the parameter $\lambda$. (b) Theoretical dependence of the uncertainty in the calculated exchange fraction, $\sigma_{\varepsilon_{obs}}$, on the true exchange fraction $\varepsilon$ for probe-exit (solid line) and probe-entry (dashed line) experiments, derived from error propagation analysis. (c, d) Model predictions for the coupled probability distributions $P^{2\phi}(\varepsilon_{obs})$ and $P^{1\phi}(\varepsilon_{obs})$ (green and pink, Eqs. 5a and 5b, respectively) for probe-exit (c) and probe-entry (d) experiments, calculated for different values of the fluorescence uncertainty $\sigma_F$ and fixed $\varepsilon^* = 0.5$ and $\lambda = 1$.

#### S5.2 Deriving the uncertainty in exchange fraction calculated from fluorescence intensity.

We calculate the fraction of exchanged outer leaflet lipids, $\varepsilon_{obs}$, of an aGUV as follows:

$$\varepsilon_{exit} = 2\left(1 - \frac{F_A}{\bar{F}_S}\right), \qquad S1a$$

$$\varepsilon_{entry} = 2\frac{F_A}{\bar{F}_S}, \qquad S1b$$

where Eq. S1a or S1b are appropriate for a probe-exit or probe-entry experiment, respectively. In Eqs. S1, $F_A$ is the average equatorial fluorescence intensity of an individual aGUV and $\bar{F}_S$ is the average fluorescence intensity of $N_S$ symmetric control GUVs at a probe concentration representing complete exchange of both leaflets:

$$\bar{F}_S = \frac{1}{N_S}\sum_{i=1}^{N_S} F_{S,i}\,. \qquad S2$$

$\bar{F}_S$ is thus a reference intensity for calculating outer leaflet exchange; for either mode of probe transfer, the intensity of an asymmetric GUV at 100% outer leaflet exchange is half the value of $\bar{F}_S$. Defining $f = F_A/\bar{F}_S$, we rewrite Eqs. S1:

$$\varepsilon_{exit} = 2 - 2f, \qquad S3a$$

$$\varepsilon_{entry} = 2f. \qquad S3b$$

Using standard rules for error propagation, the uncertainty in $f$ is:

$$\sigma_f = f\sqrt{\left(\frac{\sigma_{F_A}}{F_A}\right)^2 + \left(\frac{\sigma_{\bar{F}_S}}{\bar{F}_S}\right)^2}. \qquad S4$$

Rearranging Eqs. S3 to solve for $f$, we have:

$$f_{exit} = 1 - \varepsilon_{exit}/2, \qquad S5a$$

$$f_{entry} = \varepsilon_{entry}/2. \qquad S5b$$

Inserting this result into Eq. S4 gives:

$$\sigma_{f_{exit}} = (1 - \varepsilon_{exit}/2)\sqrt{\left(\frac{\sigma_{F_A}}{F_A}\right)^2 + \left(\frac{\sigma_{\bar{F}_S}}{\bar{F}_S}\right)^2}, \qquad S6a$$

$$\sigma_{f_{entry}} = \varepsilon_{entry}/2\sqrt{\left(\frac{\sigma_{F_A}}{F_A}\right)^2 + \left(\frac{\sigma_{\bar{F}_S}}{\bar{F}_S}\right)^2}. \qquad S6b$$

Applying the rules for error propagation to Eqs. S3, we have:

$$\sigma_{\varepsilon_{exit}} = 2\sigma_{f_{exit}} = (2 - \varepsilon_{exit})\sqrt{\left(\frac{\sigma_{F_A}}{F_A}\right)^2 + \left(\frac{\sigma_{\bar{F}_S}}{\bar{F}_S}\right)^2}, \qquad S7a$$

$$\sigma_{\varepsilon_{entry}} = 2\sigma_{f_{entry}} = \varepsilon_{entry}\sqrt{\left(\frac{\sigma_{F_A}}{F_A}\right)^2 + \left(\frac{\sigma_{\bar{F}_S}}{\bar{F}_S}\right)^2}. \qquad S7b$$

The expression under the radical symbol represents a propagation of the relative error inherent to measuring the fluorescence intensity of a *single* aGUV (the first term) with the uncertainty in the *average* intensity of a population of symmetric GUVs (the second term). In other words, the magnitude of the second term can be made arbitrarily small if enough sGUVs are measured. In our experiments, we typically measured 50 symmetric vesicles to determine $\bar{F}_S$. The average relative uncertainty in fluorescence of an individual GUV, $\sigma_{F_S}/F_S$, was about 15% (Tables S1-S3). The relative uncertainty in the *mean* intensity of 50 GUVs is then:

$$\frac{\sigma_{\bar{F}_S}}{\bar{F}_S} = \frac{1}{\sqrt{N_S}} \frac{\sigma_{F_S}}{F_S} \approx \frac{0.15}{\sqrt{50}} = 0.021.$$

If we assume that the relative uncertainty in the intensity of an individual aGUV is similar to that of an individual sGUV (i.e., $\sigma_{F_A}/F_A \approx \sigma_{F_S}/F_S = 0.15$), then the expression under the radical in Eqs. S7 is:

$$\sqrt{\left(\frac{\sigma_{F_A}}{F_A}\right)^2 + \left(\frac{\sigma_{\bar{F}_S}}{\bar{F}_S}\right)^2} \approx \sqrt{0.15^2 + 0.021^2} = 0.151,$$

i.e., the propagated error is dominated by the first term in the radical. We can then simplify Eqs. S7 to arrive at the final result:

$$\sigma_{\varepsilon_{exit}} \approx (2 - \varepsilon_{exit}) \frac{\sigma_{F_A}}{F_A}, \qquad S8a$$

$$\sigma_{\varepsilon_{entry}} \approx \varepsilon_{entry} \frac{\sigma_{F_A}}{F_A}. \qquad S8b$$

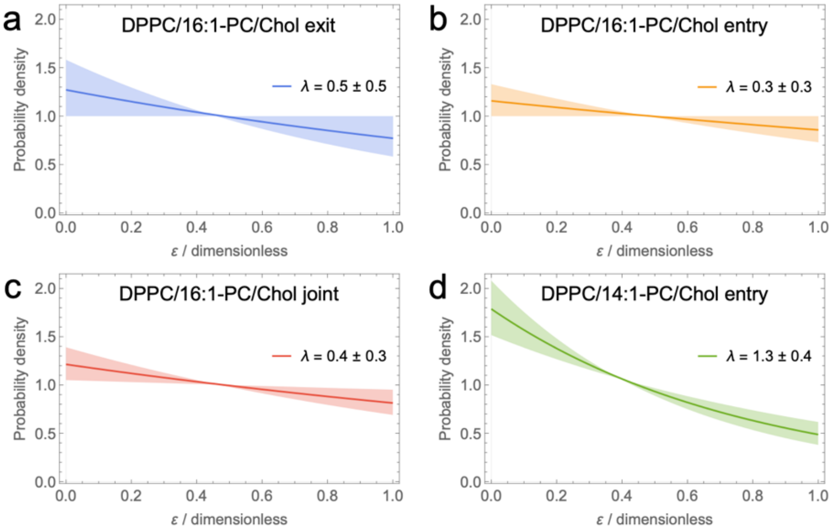


**Figure S9.** Inferred true exchange distributions $f(\varepsilon;\lambda)$ obtained from the coupled-distributions model (Eq. 3). Solid curves show the best-fit exchange distributions inferred from the experimental data, while shaded regions indicate the sensitivity of $f(\varepsilon;\lambda)$ to uncertainty in the fitted parameter $\lambda$.

### S6. Supplementary control studies

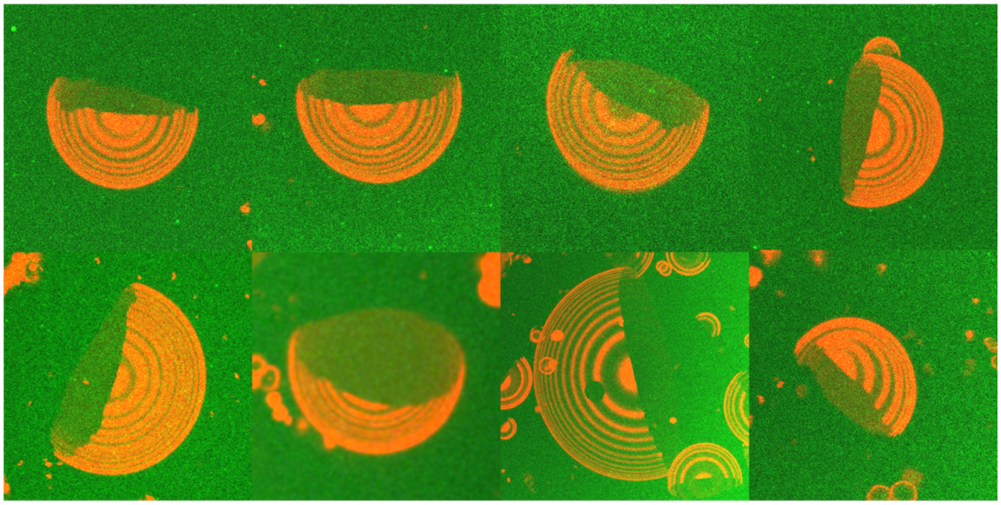

**Figure S10**. Confocal $z$-stack images demonstrating that equatorial Ld phase fractions accurately represent overall vesicle phase fractions for GUVs resting on an SLB, confirming the validity of equatorial-plane analysis.

**Table S7.** Symmetric DPPC/14:1-PC/Chol GUVs for locating ternary phase boundary.

| χDPPC | χ14:1-PC | χChol | $N_{uni}$ | $N_{2\phi}$ | $f_{2\phi}$ |
|---|---|---|---|---|---|
| 0 | 79 | 21 | 56 | 0 | 0 |
| 5 | 75 | 21 | 49 | 1 | 0.02 |
| 10 | 69 | 21 | 28 | 14 | 0.33 |
| 15 | 64 | 21 | 0 | 35 | 1 |

**Table S8.** Symmetric DPPC/16:1-PC/Chol GUVs for locating ternary phase boundary.

| χDPPC | χ16:1-PC | χChol | $N_{uni}$ | $N_{2\phi}$ | $f_{2\phi}$ |
|---|---|---|---|---|---|
| 0 | 79 | 21 | 88 | 0 | 0 |
| 6 | 73 | 21 | 96 | 0 | 0 |
| 11 | 68 | 21 | 69 | 0 | 0 |
| 15 | 64 | 21 | 5 | 83 | 0.94 |
